# LaeA-dependent production of small molecules of *Aspergillus niger* that compete with specific antibodies that bind to human immune receptors

**DOI:** 10.1101/2022.04.26.489527

**Authors:** N. Escobar, E. M. Keizer, J. F. van Neer, M. Arentshorst, J. A. G. van Strijp, P. J. A. Haas, A. F. J. Ram, P. J. Punt, H. A. B. Wösten, H. de Cock

**Affiliations:** Microbiology & Institute of Biomembranes, Department of Biology, Utrecht University, The Netherlands; Leiden University, Institute of Biology Leiden, Molecular Microbiology and Biotechnology, The Netherlands; Department of Medical Microbiology, University Medical Center Utrecht, The Netherlands; Ginkgo Bioworks, Inc., Utrecht, The Netherlands

**Keywords:** Aspergillus, immune receptors, secondary metabolites, immune evasion

## Abstract

Microorganisms secrete a variety of compounds into their environment such as proteins, carbohydrates, and secondary metabolites. These molecules play diverse roles in the interaction of microbes with their abiotic and biotic environment. Little is known about secreted fungal molecules mediating immune evasion. Here we screened culture media of three Aspergilli to assess whether these fungi secrete molecules that can compete with specific antibodies that bind to human immune receptors. Culture media of *Aspergillus fumigatus* Af293, *Aspergillus tubingensis* CBS 133792 and the non-acidifying mutant strain *Aspergillus niger* D15#26 contained components that showed competition for binding to a total of 13 receptors, of which PSGL-1, CXCR1, and CXCR2, were shared between the three species. Filtration experiments showed that most, if not all, interacting components were ≤ 3 kDa. Production of the components that competed with antibodies to bind to CD88 and CXCR2 was shown to be regulated by LaeA. The component(s) that competed for binding to CXCR1 was not only produced in the non-acidifying strain *Aspergillus niger* D15#26 but also in the non-acidifying *oahA* deletion strain of *Aspergillus niger*. Together, these data show that *Aspergillus* species might produce small molecules that interact with human immune receptors.

## Introduction

Microbial compounds known as pathogen-associated molecular patterns (PAMPs) induce a host innate immune response by binding to pattern recognition receptors. Dendritic cells (DCs), macrophages, neutrophils (PMNs), natural killers (NKs), and monocytes express a large repertoire of soluble and membrane-bound receptors (also called CD receptors) that in interaction with their ligand initiate the immune response (1–4). These immune cells are for example active on the lung epithelial layer and adjacent tissue to remove micro-organisms like fungal propagules (5,6). Cell wall molecules (e.g. chitin, mannan, glucan) are the best described fungal PAMPs. For instance, the polysaccharide β-1,3 glucan was shown to interact with complement receptor 3, dectin-1, EphA2, and ficolin-2. These interactions trigger a variety of responses like ROS production, neutrophil migration, and production of cytokines (7–9).

Microbes secrete molecules that mediate immune evasion. Various *Staphylococcus aureus* proteins involved in immune evasion have been identified by screening culture media in a competition binding assay using immune receptors (10). For example, the secreted SSLs family proteins of S. *aureus* were shown to block the interaction between PSGL-1 and P-selectin inhibiting neutrophil (PMN) recruitment. This human pathogen also secretes a virulence factor described as chemotaxis inhibitory protein (CHIPS) that inhibits PMN responses by binding to the formyl peptide receptor (FPR) and the C5a receptor (C5aR; also known as CD88). Secreted *A. fumigatus* proteases such as aspartic protease (Pep1p), metalloprotease (Mep1p), and alkaline protease (Alp1) have also been described to mediate immune evasion (11,12). Possibly, opportunistic fungal pathogens such as *A. niger, A. fumigatus*, and *A. tubingensis* secrete such compounds as part of the large variety of molecules they release into their environment. The secreted compounds of *Aspergillus* includes enzymes, secondary metabolites, sugars and organic acids (13–17). The number and type of secreted compounds vary depending on the culture conditions (18–21). By combining *in silico* predictions and shotgun proteomics it was estimated that *A. niger* can secrete over 200 proteins (22). These proteins are predicted to serve functions in cellular communication, immunity and pathogenesis, degradation of substrates, morphogenesis, and cell proliferation (13). A similar number of secondary metabolites (i.e. 145) have been isolated and / or detected from cultures of black aspergilli (*Aspergillus* section *Nigri*) (23). Secondary metabolites are low molecular weight molecules with various functions. Some metabolites are mycotoxins (e.g. fumonisin B2, ochratoxin A) causing food spoilage and being a threat for human health. Yet, for many fungal secondary metabolites a biological role has not been elucidated.

The *prtT* and *laeA* genes of *A. niger* have been shown to play a crucial role in the production of extracellular compounds (24–27). Strains with *prtT* mutations exhibit decreased extracellular protease activity when compared to the parental strain *A. niger* AB4.1 (27). This gene encodes a Zn(II)2Cys6 transcription factor and controls the expression of 6 secreted proteases including Aspergillopepsin A (PepA) and Aspergillopepsin B (PepB) (27). Transcriptional and proteomic analysis of an *A. fumigatus prtT* mutant indicated that PrtT also regulates genes involved in iron uptake, ergosterol biosynthesis, and production of secondary metabolites (28,29). Gene *laeA* encodes a putative methyltransferase domain protein and is a global regulator of secondary metabolite gene clusters in *Aspergillus*. Amongst others, synthesis of gliotoxin, sterigmatocystin, penicillin, and lovastatin is being regulated by LaeA (26). Notably, secondary metabolite production of the *A. niger* Δ*laeA* strain differed when compared to the wild-type, with both an increase and decrease in secondary metabolite levels (30). This indicates that this protein can also act as a repressor of secondary metabolite genes. Deletion of *laeA* also leads to a decrease in acidification of the culture medium (30), which is due to reduced production of citric acid, gluconic acid and / or oxalic acid. Production of these organic acids is pH dependent (31). For example, optimal production of citric acid in *A. niger* occurs at pH 2 (32), while oxalic acid production is optimal between pH 5 to 8 (33).

The global regulator LaeA was proposed to regulate secondary metabolite synthesis via chromatin remodelling and regulates amongst others secondary metabolites produced via polyketide synthases (PKS) or non-ribosomal peptide synthases (NRPS) (34). Both of these synthesis pathways require post-translational modification for activation. The 4’phosphopantetheinyl transferase (PPTase) activates PKS and NRPS. Deletion of this PPTase leads to a defect in secondary metabolite production in *A. fumigatus, A. nidulans* and *A. niger* (35–38). Other fungal secondary metabolite pathways are the mevalonate or shikimic acid pathway, which are involved in the production of terpenes or aromatic secondary metabolites, respectively (39,40).

Identification of secreted immune-reactive molecules might unravel new immune evasion strategies and may reveal novel therapeutic agents. In this study we show that culture media of *A. niger, A. fumigatus*, and *A. tubingensis* contain thermostable molecules ≤ 3 KDa that might compete with monoclonal antibodies in their interaction to receptors involved in immune recognition. It is also shown that LaeA of *A. niger* is a repressor of the production of the compounds that compete with binding to CD88 and CXCR2, while secretion of the compound binding to CXCR1 is increased under non-acidifying medium conditions.

## Materials and methods

### Strains, growth conditions, isolation of culture medium, and extracellular fractions

Strains (Table 1, Figure 1) were grown for 3 days at 37 °C in minimal medium (MM; 6 g L^-1^ NaNO_3_, 1.5 g L^-1^ KH_2_PO_4_, 0.5 g L^-1^ KCl, 0.5 g L^-1^ MgSO_4_.7H_2_O, 0.2 mL L^-1^ Vishniac; pH 6.0) supplemented with 25 mM glucose and 1.5 % agar to obtain conidia. Conidia were harvested with 0.005 % (v/v) Tween-80 in 0.85 % (w/v) NaCl. A total number of 10^10^ conidia was used to inoculate 250 mL transformation medium (TM; MM supplemented with 5 g L^-1^ yeast extract (Becton, Dickinson and Company, Le-Pont-De-Claix, France) and 2 g L^-1^ casamino acids (Becton, Dickinson and Company, Le-Pont-De-Claix, France)) in a 1 L Erlenmeyer. Cultures were grown for 16 h at 30 °C and 250 rpm in MM with 25 mM xylose or maltose as carbon source. Mycelium was harvested by filtration over 3 layers of Miracloth (Merck Millipore, Darmstadt, Germany) and washed with 50 mL PBS (137 mM NaCl, 2.7 mM KCl, 3.8 mM Na_2_HPO_4_·2H_2_O, 1.5 mM KH_2_PO_4_). 10 g wet weight mycelium was transferred to a 500 mL Erlenmeyer containing 150 mL MM supplemented with either 25 mM xylose or maltose. Cultures were grown for 3 days at 30 °C and 250 rpm. Pellet formation was followed using light microscopy (Axioskop 2 plus, Carl Zeiss) and their surface area was measured after 72 h using Image J (https://imagej.nih.gov/ij/). Mycelium and mycelial fragments were removed by filtering over 3 layers of Miracloth and a 0.22 μm filter (Carl Roth GmbH + co, KG, Karlsruhe, Germany). Xylose- and maltose-culture media were mixed 1:1 for analysis. In order to obtain small secreted molecules, supernatants were filtered using a 3 kDa cut-off Amicon ultra centrifugal filter (Merck Millipore, Darmstadt, Germany). Culture media and ≤ 3 KDa fractions were lyophilized and suspended in PBS. Strains lacking the *pptA* gene were grown in MM medium with either xylose or maltose as carbon source, which was mixed 1:1 with siderophore medium and 10 mM lysine (Sigma Aldrich, St. Louis, France). Siderophore medium was made by growing *A. niger* N402 in liquid MM in the absence of Fe^++^, with 5 mM glutamine (Sigma Aldrich, St. Louis, France) as a nitrogen source and 50 mM glucose as carbon source. *A. niger* was cultured for a total of 48 h at 200 rpm and after the first 24 hours of growth, fresh glutamine (5 mM) was added. Mycelium was removed from the medium by filtering over 3 layers of Miracloth, the pH was set to 6.0 and the siderophore culture medium was autoclaved (37).

**Table 1.** Strains used in this study.

| Strain name | Parent | Relevant genotype | Reference |
| --- | --- | --- | --- |
| *A. niger* |  |  |  |
| N402 |  | Short conidiophore mutant of NRRL3 | (42) |
| D15#26 | N402 | An UV-generated *pyrG<sup>-</sup> prtT<sup>-</sup>, laeA<sup>-</sup>* strain isolated as a non-acidifying mutant (*ac<sup>-</sup>*) | (30) |
| JN21.1 | D15#26 | pAO4_13 | (30) |
| JN22.7 | D15#26 | pAO4_13-*laeA* | (30) |
| JN23.1 | AB4.1 | pAB4.1 *pyrG<sup>+</sup>* |  |
| JN24.6 | AB4.1 | $\Delta laeA::AopyrG$ | (30) |
| AB1.13 | AB4.1 | *pyrG<sup>-</sup> prtT<sup>-</sup>* | (66) |
| AB1.13$\Delta oahA\#76$ | AB1.13 | $\Delta oahA::pyrG$ | (67) |
| JP1.1 | N402 | $\Delta pptA::AopyrG$ | (36) |
| MA870.1 | JN24.6 | $\Delta laeA$, $\Delta pptA::hygR$ | This publication |
| AF11#56 | AB4.1 | $\Delta prtT::pyrG$ | (68) |
| *A. fumigatus* |  |  |  |
| Af293 |  | Clinical isolate from lung tissue | (46) |
| *A. tubingensis* |  |  |  |
| CBS 133792 |  | Clinical isolate from an immuno-compromised patient suffering from osteomyelitis | (48) |

**Figure 1.**
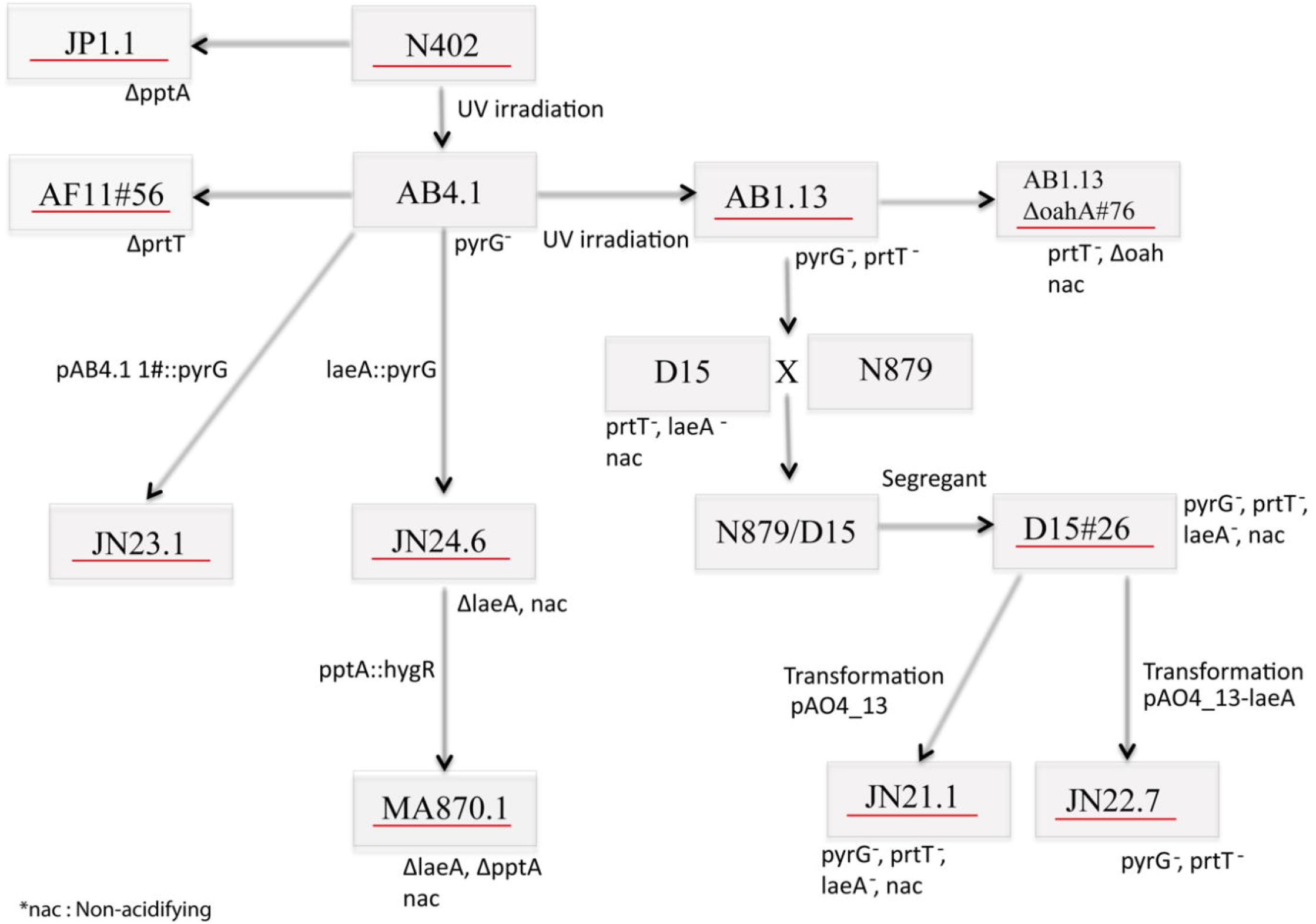
Schematic overview of strains used in this study (underlined in red). Adapted from (30). nac: non acidifying.

### Transformation *A. niger*

The *pptA* gene was deleted in strain JN24.6 (*laeA::AOpyrG*) (30) using the split marker method, with hygromycin as selection marker (41). Flanks of *pptA* were PCR amplified using Phire Hot Start II DNA Polymerase (Thermo Scientific), with genomic DNA of N402 (42) as template and primers pptAP1f and pptAP2r and primers pptAP3f and pptAP4r to give the 5’ pptA flank of 857 bp and the 3’ pptA, flank of 862 bp, respectively. The hygromycin fragments were PCR amplified using plasmid pAN7.1 (43) as template and primers hygP6f and hygP9r to give the 5’ hygR fragment (1794 bp) and primers hygP8f and hygP7r to give the 3’ hygR fragment (1644 bp). Split marker fragments were obtained using fusion PCR (5’ pptA-hygR (2633 bp) and 3’pptA-hygR, (2485 bp)) and column-purified before further use, using the GeneJET Gel Extraction Kit (Thermo Scientific). All primers that were used are listed in (Supplementary Table 1). The split marker fragments were transformed to strain JN24.6 and transformants were purified twice on complete medium (CM; MM supplemented with 2 g L^-1^ tryptone, 1 g L^-1^ casamino acids, 1 g L^-1^ yeast extract, 0.5 g L^-1^ yeast ribonucleic acids, pH 6.0) with siderophore medium, 10 mM lysine and 100 μg/mL hygromycin, resulting in strain MA870.1 (*pptA::hygR, laeA::AOpyrG*). Correct deletion of *pptA* was confirmed by diagnostic PCR (Supplementary Figure 1), clone 1 was used for experiments described.

### SDS PAGE

Proteins from culture media were precipitated with 4 volumes of acetone (Merck, Darmstadt, Germany) for 16 h at 20 °C. Samples were centrifuged at 10,000 g for 15 min, re-suspended in 2 x SDS sample buffer (20 % glycerol (LPS Benelux, The Netherlands), 4 % SDS (JT Baker, Deventer, The Netherlands), 100 mM Tris-HCl pH 6.8 (Roche, Mannheim, Germany), 0.01 % bromophenol blue (Acros Organics, Geel, Belgium) and 5 %β-mercaptoethanol (Sigma-Aldrich, St. Louis, France)), and heated for 10 min at 100 °C. Samples and Low Molecular Weight Marker (14.000-70.000 Da) (Sigma-Aldrich, St. Louis, France) were loaded on 12 % SDS-PAA gels and stained with 0.1 % Coomassie Brilliant Blue G250 (Sigma-Aldrich, St. Louis, France) in 25 % methanol (Merck, Darmstadt, Germany) and 10 % acetic acid (Merck, Darmstad, Germany).

### PMNs, and PBMCs isolation and competition for receptor binding assay

Polymorphonuclear neutrophils (PMNs) and peripheral blood mononuclear cells (PBMCs) were routinely isolated from whole blood of healthy donors following the Histopaque-Ficoll gradient protocol (10). Written informed consent was obtained from all subjects according to the Declaration of Helsinki. Approval was obtained from the medical ethics committee of the University Medical Center Utrecht (Utrecht, The Netherlands). Competition for receptor binding assay (CBA) was performed using commercial phycoerythrin (PE)-, fluorescein isothiocyanate (FITC)-, and allophycocyanin (APC)-conjugated antibodies (Supplementary Table 2) as described (10) with some modifications. Briefly, 100 μL of PMNs and 150 μL PBMCs (each 10^7^ cells mL^-1^) were mixed and centrifuged for 5 min at 7,000 g. The pellet was resuspended in 1 mL of PBS mixed 1:1 with filtrated fungal culture medium supernatant at 4 °C. If necessary, pH of the samples was adjusted to 7 using 0.1 M NaOH. PBS mixed 1:1 with only the culture medium was used for control in a subset of experiments. After incubation for 15 min at 4 °C, 35 μL of the mixture was incubated with antibodies (concentrations in Supplementary Table 2) at 160 rpm for 45 min at 4 °C in 96-well U-plates (Corning, New York, USA). Cells were washed with 150 μL RPMI medium (Life Technologies, Paisley,UK) containing 0.05 % human serum albumin (Sanquin, Amsterdam, The Netherlands) and centrifuged at 1200 g for 8 min at 4 °C. Cells were fixed with 1 % paraformaldehyde (Sigma-Aldrich, Buchs, Switzerland) and fluorescence was measured by flow cytometry (FACSVerse, BD). Geometric mean fluorescence from neutrophils, monocytes, and lymphocytes was determined using FlowJo software (version V10.1, TreeStar, USA) to gate each cell population (lymphocytes, monocytes, and neutrophils). Representative FACS plots of control cells and supernatant treated cells can be found in supplementary figure 10. Reduction in fluorescence due to competition for binding with molecules within the culture media was calculated by dividing the mean signal by that of buffer-treated cells. Values were inverted and receptors with geometric means ≥ 2 were scored positive for binding of molecules within the culture medium. Reproducibility of the assay was confirmed by using biological triplicates with culture media and blood cells from independent cultures and donors, respectively. Receptors that scored positive were re-measured in ≥ 3 independent experiments, receptors that scored positive in at least 2 experiments were scored as responsive. Data was visualized using R software (https://www.r-project.org/) and boxplots were generated with the ggplot2 package (http://ggplot2.org). Data from neutrophils, monocytes, and lymphocytes were included in the same plot and used to calculate median and quartiles. Data outside the boxplot whiskers were taken as outliers.

### Characterization of D15#26 3KDa supernatant

Heat stability was assessed by incubating samples at 100 °C for 60 min. Samples were lyophilized and re-suspended in PBS. (Poly)peptides were precipitated from the culture medium for 30 min at 4 °C after adding acetone (Merck, Darmstadt, Germany) in a 1:1 ratio. After centrifugation at 11,000 g for 15 min, the pellet was air-dried, while the supernatant was dried using a rotoevaportor-RE (Büchi, Flawil, Switzerland). Fractions were resuspended in PBS and tested in CBA. Hydrophobic compounds were extracted from the culture medium by mixing with 3 volumes of ethyl acetate (Acros Organics, Geel, Belgium). Aqueous phases were collected, and treatment was repeated twice. Ethyl acetate fractions were pooled and dried with a rotoevaporator, while aqueous fractions were lyophilized. Fractions were resuspended in PBS in 1/3 of the original volume and tested in CBA.

### Purification of D15#26 3KDa supernatant

#### Sep-Pak C18 column purification

Ten C18 solid phase cartridges (Sep-pak; Waters, Milford, MA) were used to load 50 mL of culture medium that had been lyophilized and resuspended in 50 mL of PBS. Columns were eluted stepwise with 0, 10, 30 50, 70 and 100 % methanol (v/v). Fractions with the same percentage of methanol were pooled, lyophilized, resuspended in PBS, and tested in CBA. Fractions obtained from elution with 10 - 50 % methanol were pooled and concentrated 100-fold for LC-MS analysis.

#### *LC-MS* and *Preparative HPLC*

LC-MS was performed on a Shimadzu SCL-10A controller system (Shimadzu Cooperation, ‘s-Hertogenbosch, The Netherlands) coupled to a Shimadzu pump LC10-AD and a Shimadzu CTO-10AS column oven. A Reprosil-Pur C18-AQ column (Particle size = 5 μm, Pore size = 120Ǻ, 250 x 4,6 mm; Reprosil) was loaded with 50 μL 100-fold concentrated 10 - 50 % methanol pooled fraction. A 0 −100 % gradient elution (Supplementary Table 3) with water and acetonitrile (JT Baker, HPLC grade) was used for separation of molecules during 60 min. The flow rate was 1 mL min^-1^ and compounds were detected with a UV detector at 214 nm (Shimadzu SPD-10A). Mass spectrometry was done using a Finnigan LCQ Deca XP Max (Thermo Electron, Massachusetts, USA).

Preparative HPLC was run on a Shimadzu SCL-10A controller system coupled with a Shimadzu LC-8a pump and a Shimadzu SPD-10A UV detector. A Reprosil-Pur C18-AQ column (particle size =10 μm, pore size= 120Ǻ, 250 x 22 mm; brand) with a Reprosil-Pur C18-AQ guard column (particle size =10 μm, pore size= 120Ǻ, 30×22 mm) was injected with 450 μL of concentrated 10-50 % methanol pooled fraction. For sample separation a 0 to 60 % gradient elution with water and acetonitrile over 100 min was used with a flow rate of 12.5 mL min^-1^. Ninety-five 13 mL fractions were collected using a Gilson Liquid Handler 215. Fractions were pooled in equal ratios (1 mL each fraction), obtaining 16 fractions that were dried in a SpeedVac, resuspended in 1 ml of PBS, and subjected to CBA.

### Prediction of *A. tubingensis* SM clusters

*A. tubingensis* CBS 134.48 genome v 1.0 (44) (http://genome.jgi.doe.gov/Asptu1/Asptu1.download.html)was used to predict the number of genes clusters involved in biosynthesis of secondary metabolites. Analysis was performed using anti-SMASH parameters (http://antismash.secondarymetabolites.org). By using the “homologous gene cluster” tool, gene clusters were identified with similarity to gene clusters in the *A. niger* ATCC 1015 (http://genome.jgi.doe.gov/Aspni5/Aspni5.download.html) and *A. fumigatus* Af293 (http://genome.jgi.doe.gov/Aspfu1/Aspfu1.download.html) genomes.

## Results

### Characterization of fungal cultures

Protein profiles of *Aspergillus* culture media were monitored after 24, 48, and 72 h of growth. To this end, samples of maltose and xylose media were mixed in a 1:1 ratio, precipitated, and analyzed by SDS-PAGE (Figure 2). Protein profiles showed a high variation between the different strains. *A. niger* N402 showed bands > 68 kDa that were reduced in intensity in *A. niger* D15#26. *A. tubingensis* that also belongs to the *Aspergillus* section *Nigri* showed a protein profile different from the two *A. niger* strains. *A. fumigatus* showed also a distinct profile containing high (55-80 kDa) and low molecular weight bands (15-30 kDa). The *A. niger* D15#26 strain does not acidify the culture medium. Indeed, pH of the culture medium had increased to 7 after 72 h of growth (Table 2). In contrast, *A. niger* N402 and *A. tubingensis* had lowered the pH to 5.5, while the pH of the culture medium of *A. fumigatus* had increased to pH 8. All cultures showed pelleted growth, but morphology was different (Table 2). The *A. niger* strain D15#26 produced smaller pellets mixed with dispersed growth when compared to wild-type *A. niger* N402 (Supplementary Figure 2).

**Figure 2.**
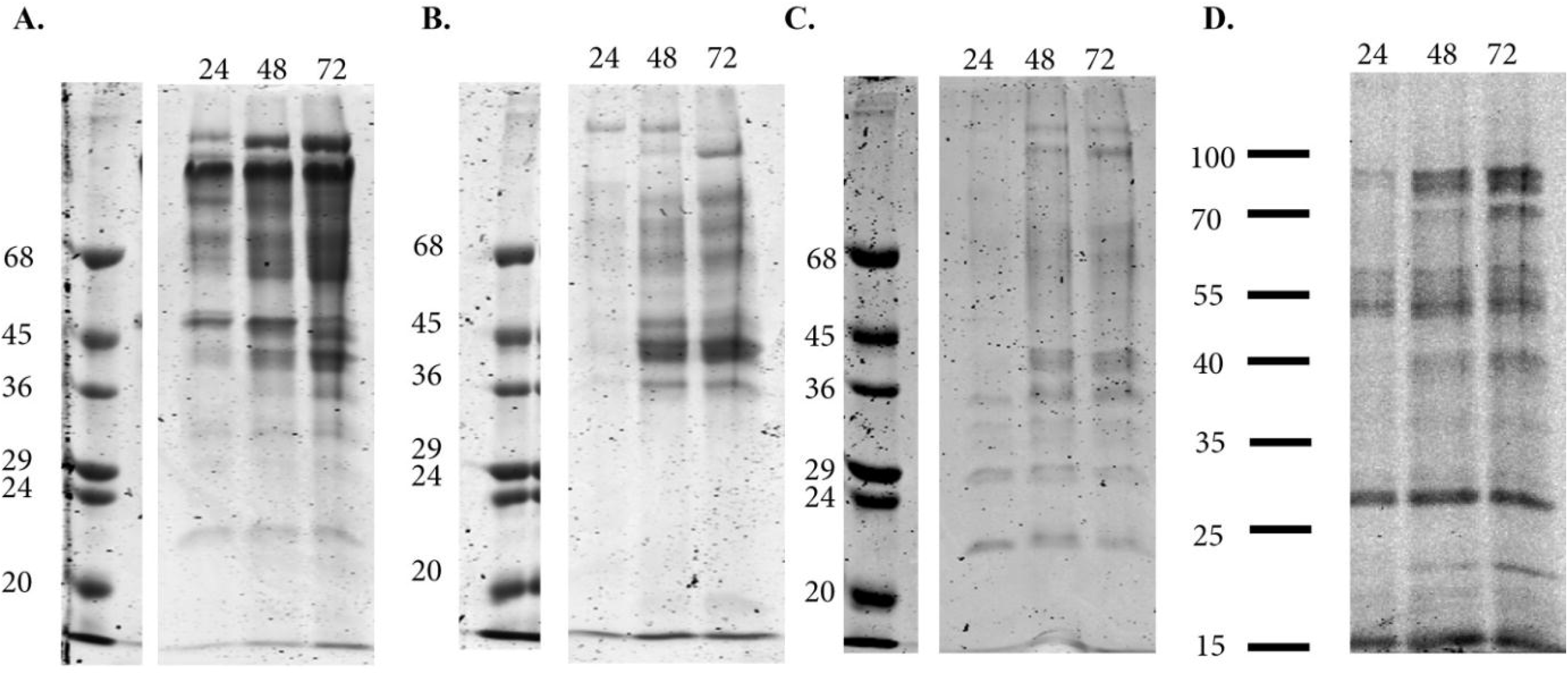
CBB stained SDS PAA gels of mixed maltose and xylose culture media of *A. niger* N402 (A), *A. niger* D15#26 (B), *A. tubingensis* (C), and *A. fumigatus* 293 (D) after 24, 48, and 72 h of growth.

**Table 2.** pH of *Aspergillus* culture media after 24, 48, and 72 h of growth and mycelium morphology after 72 h. The pH of culture medium at 0 h was 6.

| Strain | pH |  |  | Growth characteristic after 72 h of growth |
| --- | --- | --- | --- | --- |
|  | 24 h | 48 h | 72 h |  |
| *A. niger* N402 | 4.5 | 5.5 | 5.5 | Dense pellets ($\pm 450,000 \mu\text{m}^2$) |
| *A. niger* D15#26 | 5.5 | 6.5 | 7 | Pellets ($\pm 50,000 \mu\text{m}^2$) with some dispersed growth |
| *A. tubingensis* | 4.5 | 5 | 5.5 | Dense pellets ($\pm 230,000 \mu\text{m}^2$) |
| *A. fumigatus* 293 | 7 | 7 | 8 | Pellets ($\pm 22,500 \mu\text{m}^2$) with dispersed growth |

Mixed maltose and xylose culture media were used to challenge PMNs and PMBCs in a competition binding assay (CBA) in order to detect secreted compounds that interact with human immune receptors. A component(s) in the culture medium of *A. niger* N402 (Figure 3A) competed with monoclonal antibodies directed against PSGL-1 in several of the experiments but variation was too high to reach a median ≥ 2 when compared to the control. Possibly, the lack of response with other receptors was due to extracellular proteases degrading binding proteins. We therefore used a derivative of N402, strain D15#26, that has low protease activity due to a *prtT* mutation. Culture media of *A. niger* D15#26 indeed contained a component(s) that competed for binding to CD141, while receptors PSGL-1, CXCR1, CXCR2, and CD47 were competing for binding in several experiments but had a median < 2 (Figure 3B). Anion / cation chromatography was performed as a first step for protein purification and fractions were tested in CBA. However, competition for binding to receptor molecules was not observed in any of the fractions (Supplementary Figure 3). The purification procedure included a dialysis step with membranes with a 12 kDa cut off. To assess whether small molecular weight molecules were responsible for the CBA response culture media were filtered using 3 kDa filters and fractions were tested with a subset of receptors. Competing for binding responses of the ≤ 3 kDa fraction of D15#26 were stronger when compared to whole culture medium (see boxplot median Figure 4A). PSGL-1, CXCR1, and CXCR2 showed a strong response, while response of CD88 was also close to 2-fold. The N402 ≤ 3 kDa fraction resulted in signals of receptors PSGL-1, CXCR1, and CXCR2 just below the 2-fold response threshold (Figure 4B). CBA was also performed with the complete set of receptors using whole culture media (Figure 5) and ≤ 3 kDa fractions (Figure 6) of clinical *A. tubengensis* and *A. fumigatus* strains. Culture media of these strains contained components that competed with binding of a total of 11 receptors, of which PSGL-1, CXCR1, CXCR2, CD192, CD47, CD13, and CD99 were shared between both clinical isolates. In most cases activity was present in the ≤ 3 kDa fractions (Figure 6).

**Figure 3.**
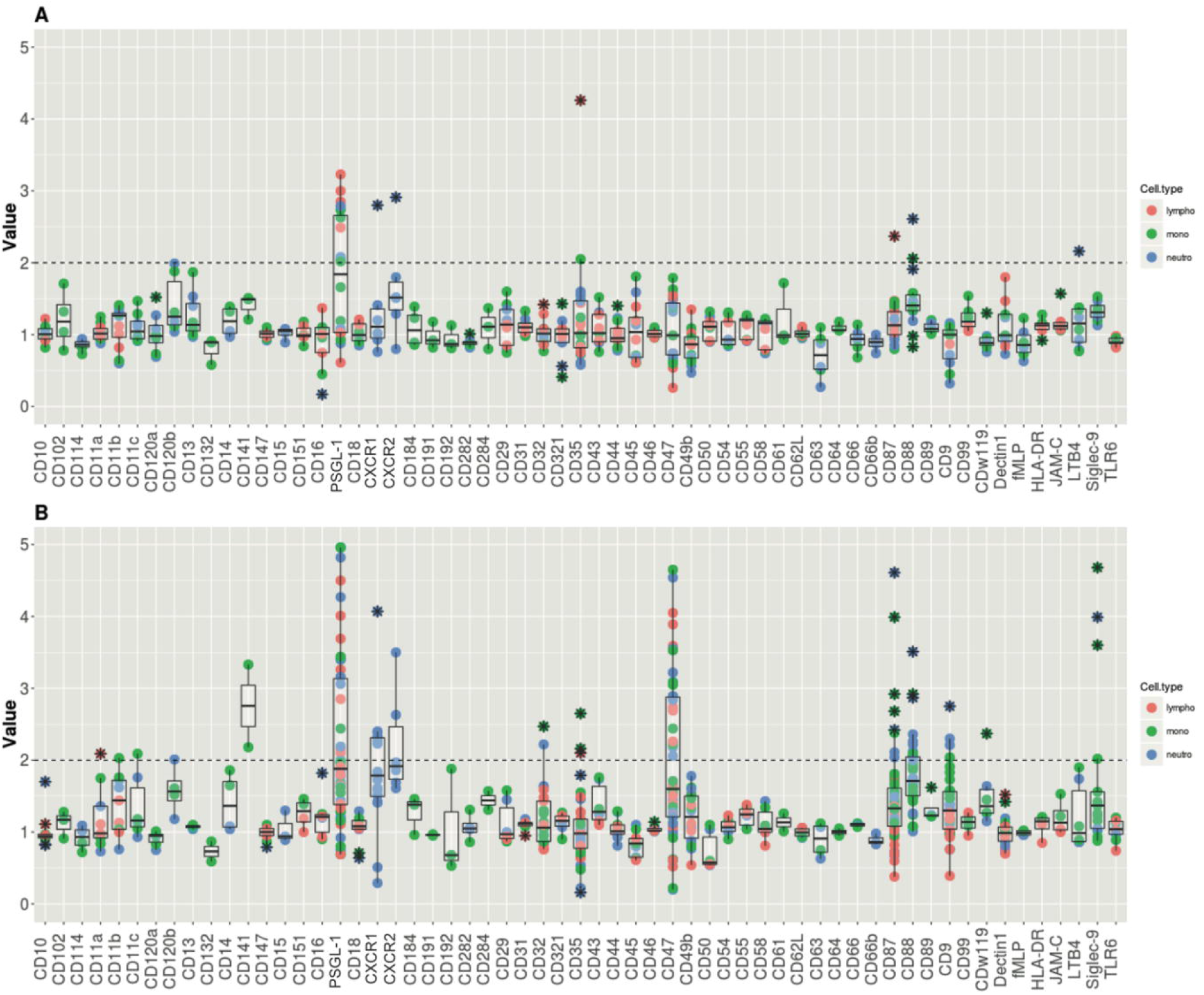
Competition binding assay of culture media of *A. niger* N402 (A) and *A. niger* D15#26 (B). Lymphocytes, monocytes, and neutrophils are represented with red, green, and blue dots, respectively, * represents outliers. Y-axis represent the inverted geometric mean of fluorescence, the X-axis represent the used receptors in the CBA, data points above the dotted line (2) are scored as positive for binding of molecules from the culture medium. Samples were tested in 3 independent experiments.

**Figure 4:**
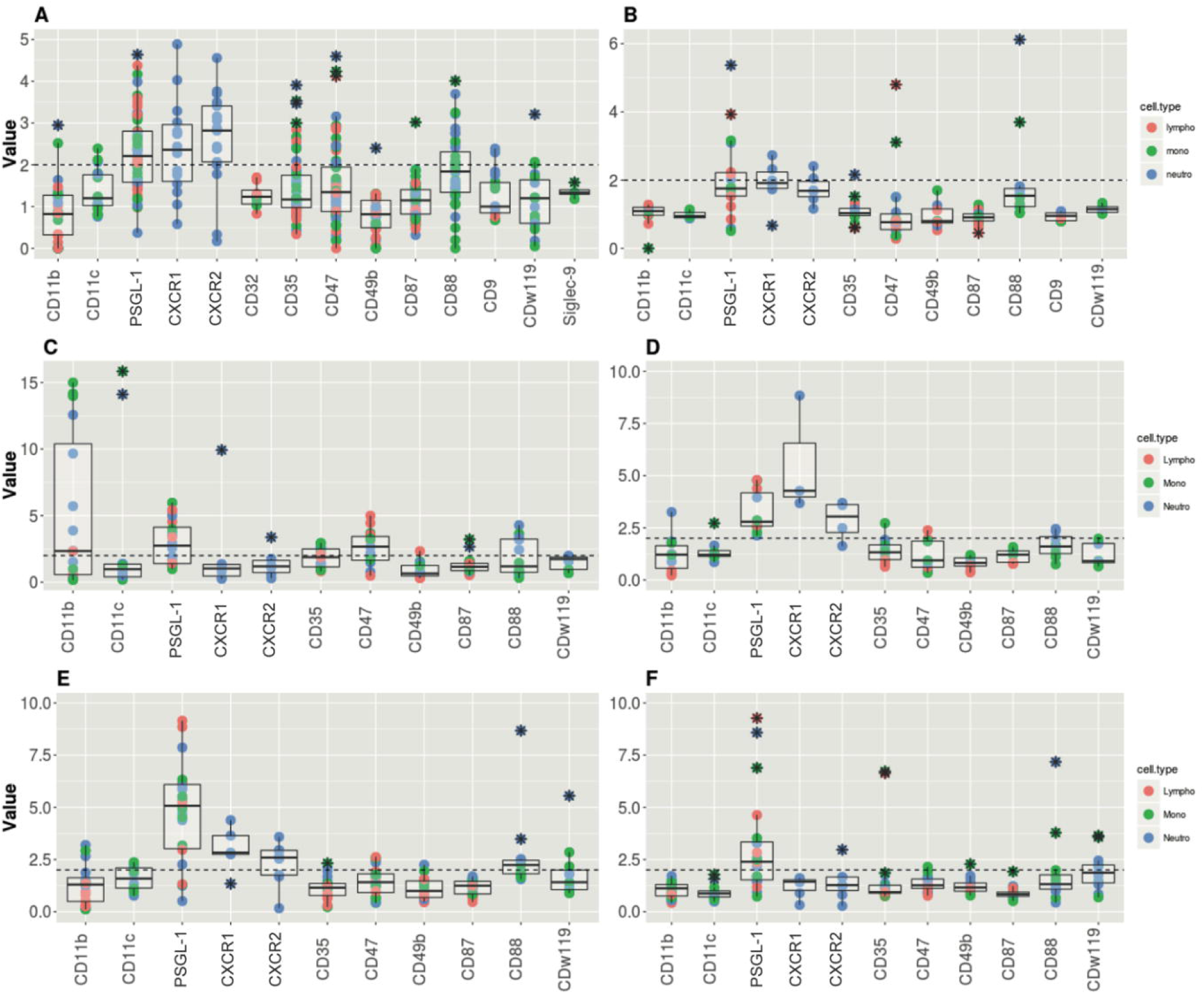
Competition binding assay using ≤ 3kDa fractions of culture media of D15#26 (A), N402 (B), JN22.7 (C), JN21.1 (D), JN24.6 (E), and JN23.1 (F). Lymphocytes, monocytes, and neutrophils are represented with red, green, and blue dots, respectively, * represents outliers. Y-axis represent the inverted geometric mean of fluorescence, the X-axis represent the used receptors in the CBA, data points above the dotted line (2) are scored as positive for binding of molecules from the culture medium. Samples were tested in 3 independent experiments.

**Figure 5.**
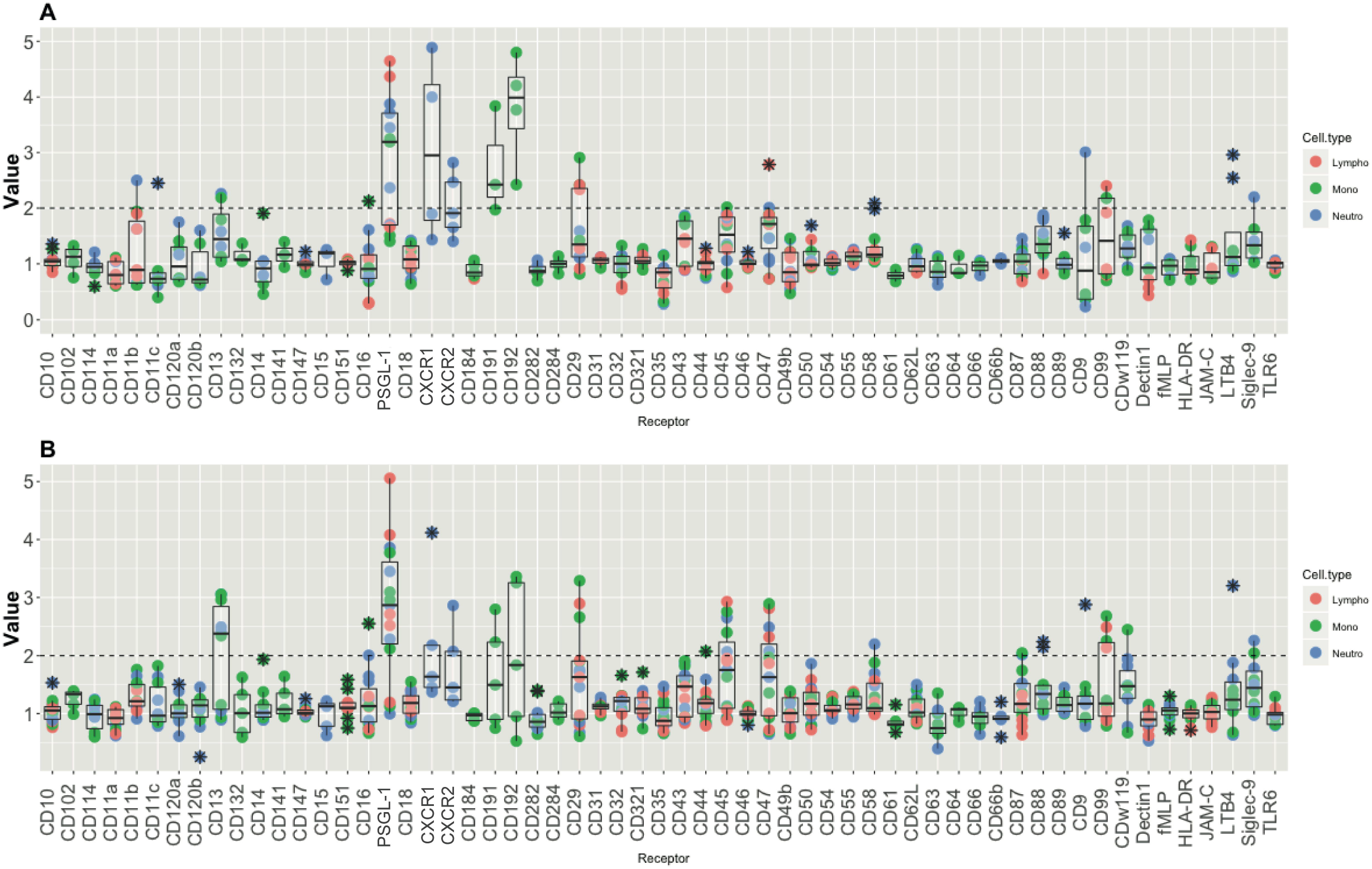
Competition binding assay of *A. fumigatus* Af293 (A) and *A. tubingensis* (B) whole culture media. Lymphocytes, monocytes, and neutrophils are represented with red, green, and blue dots, respectively, * represent outliers. Y-axis represent the inverted geometric mean of fluorescence, the X-axis represent the used receptors in the CBA, data points above the dotted line (2) are scored as positive for binding of molecules from the culture medium. Samples were tested in 3 independent experiments.

**Figure 6.**
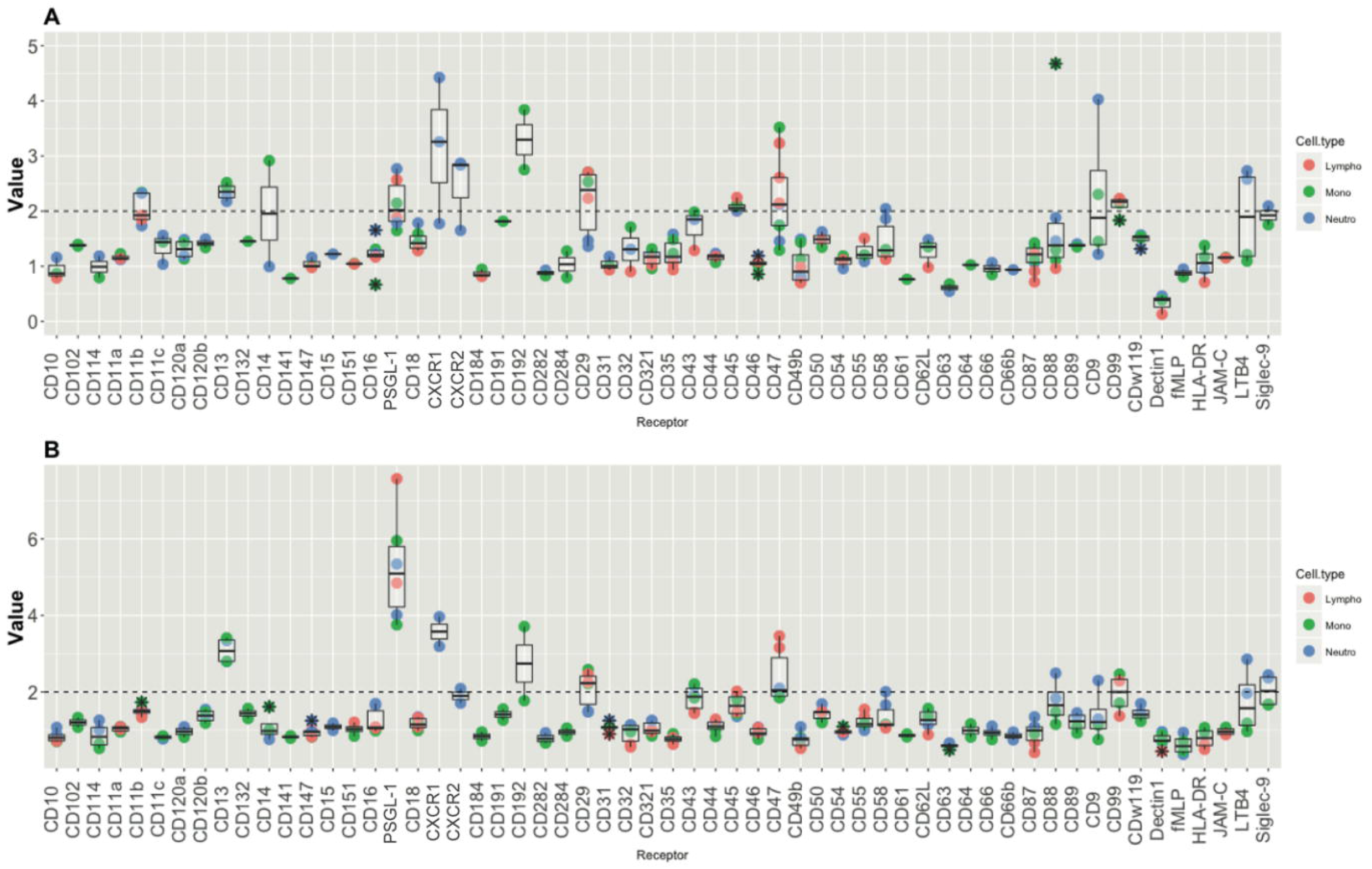
Competition binding assay of *A. fumigatus* Af293 (A) and *A. tubingensis* (B) ≤ 3 kDa fractions. Lymphocytes, monocytes, and neutrophils are represented with red, green, and blue dots, respectively, * represent outliers. Y-axis represent the inverted geometric mean of fluorescence, the X-axis represent the used receptors in the CBA, data points above the dotted line (2) are scored as positive for binding of molecules from the culture medium. Samples were tested in 3 independent experiments.

### Characterization D15#26 ≤ 3 kDa fraction

Incubation at 100 °C for 60 min did not affect competition activity of molecules within the ≤ 3 kDa fraction of D15#26 to receptors PSGL-1, CXCR1, CXCR2, and CD88 (Supplementary Figure 4). Activity was also not reduced by removing proteins by precipitation with acetone. In agreement, competition activity was absent in the protein fraction. Extraction with ethyl acetate also did not affect activity in the aqueous phase (Supplementary Figure 5). These results suggest that competing molecules are hydrophilic. Next, the ≤ 3 kDa fraction of D15#26 was loaded onto a C18 Sep-pak column and molecules were eluted using a methanol gradient. Both flow-through as well as fractions eluted with 10 - 50 % methanol contained components with competition activity to receptors CXCR1, CXCR2 and CD88, competing activity to the PSGL-1 receptor was found in all fractions (Supplementary Figure 6). Analysis of the D15#26 ≤ 3 kDa fraction by LC-MS indicated that it contained around 30 peaks (Supplementary Figure 7). CBA of the fractions collected from the LC-MS analysis was performed using the same subset of receptors used to test the ≤ 3 kDa fractions. Competing activity was not found in the tested pooled fractions. Loss of activity could be explained due to a partial binding of active components to C18-AQ column and therefore a decrease of the sample concentration or the inability of the hydrophilic compounds to bind to the C18-AQ column.

### LaeA is involved in production of CXCR1, CXCR2, and CD88 binding compounds

D15#26 is a strain resulting from UV mutagenesis (Figure 1) carrying mutations in *pyrG, prtT*, and *laeA* (30). Here, it was addressed whether *laeA* impacts the production of immune receptor competing molecules in *A. niger*. To this end, the ≤ 3 kDa fractions from the culture media of D15#26, a *laeA* complemented derivative of D15#26 (JN22.7), and its control (JN21.1) that only has a *pyrG* complementation (Figure 1) were tested in the CBA using a subset of receptors including PSGL-1, CXCR1, CXCR2, and CD88 that were found to be responsive with secreted molecules of D15#26 (Figure 4A, B). The ≤ 3kDa fraction of the *laeA* complemented strain was positive for CD11b, CD47, and PSGL-1, but not for receptors CD88, CXCR1, and CXCR2 (Figure 4C). The control strain JN21.1 behaved as D15#26 (Figure 4D) showing that complementation of *pyrG* did not affect production of the small competing compounds.

Gene *laeA* was previously inactivated in strain AB4.1 (Figure 1) by homologous recombination with a *pyrG* gene cassette (30). The ≤ 3 kDa fractions of the Δ*laeA* deletion strain (JN24.6) and its control (JN23.1, *pyrG* complementation of AB4.1) were tested in the CBA. JN24.6 produced competing activity with CD88, PSGL-1, CXCR1, and CXCR2 (Figure 4E), while its control strain only produced competing activity with PSGL-1 (Figure 4F). These results show that the absence of LaeA is correlated with production of CD88, CXCR1, and CXCR2 competing components. Considering that these strains contain an intact *prtT* gene, the production of competing activity with receptors CD88, PSGL-1, CXCR1, and CXCR2 is not related with *prtT* mutations.

### Medium acidification and production of immune-reactive components

Inactivation of *laeA* affects production of the organic acids citric acid, gluconic acid, and oxalic acid. Indeed, acidification was absent in the medium of 72 h-old cultures of strains lacking LaeA (Table 3). In contrast, the *laeA* complemented strain JN22.7 acidified the medium to pH 3, while pH dropped to 4-5 in the case of JN23.1 and AB4.1. Inactivation of the oxaloacetate hydrolase gene *oahA* in strain AB1.13 containing an intact *laeA* copy also results in absence of acidification of the culture medium (30). The ≤ 3 kDa fraction of the Δ*oahA* strain contained molecules with activity to PSGL-1 and CXCR1 in the CBA (Figure 7). Taken together, CXCR1 reactive components are produced by the *A niger* strain when medium is not acidified. Production of compounds binding to CD88 and CXCR2 receptor are not produced and are therefore LaeA dependent.

**Table 3.** pH of the medium of 72-h-old *A. niger* cultures grown in MM in the presence of xylose or maltose. Strains JP1.1, JN24.6 and MA870.1 were grown in siderophore medium

| Strains |  | Medium addition | pH Maltose | pH Xylose |
| --- | --- | --- | --- | --- |
| D15#26 | $\Delta\text{pyrG}, \Delta\text{prtT}, \Delta\text{laeA}$ | | 7 | 7 |
| JN21.1 | $\Delta\text{pyrG}, \Delta\text{prtT}, \Delta\text{laeA}$ | | 7 | 7 |
| JN22.7 | $\Delta\text{pyrG}, \Delta\text{prtT}$ | | 3 | 3 |
| JN23.1 |  |  | 5 | 4 |
| JN24.6 | $\Delta\text{laeA}$ | | 7 | 7 |
| AB1.13 | $\Delta\text{pyrG}, \Delta\text{prtT}$ | | 4.5 | 4.5 |
| AB1.13$\Delta\text{oahA}\#76$ | $\Delta\text{pyrG}, \Delta\text{prtT}, \Delta\text{oahA}$ | | 7 | 7 |
| JP1.1 | $\Delta\text{pptA}$ | Siderophore medium | 5 | 5.5 |
| JN24.6 | $\Delta\text{laeA}$ | Siderophore medium | 7 | 7 |
| MA870.1 | $\Delta\text{pptA}, \Delta\text{laeA}$ | Siderophore medium | 6 | 7 |

**Figure 7.**
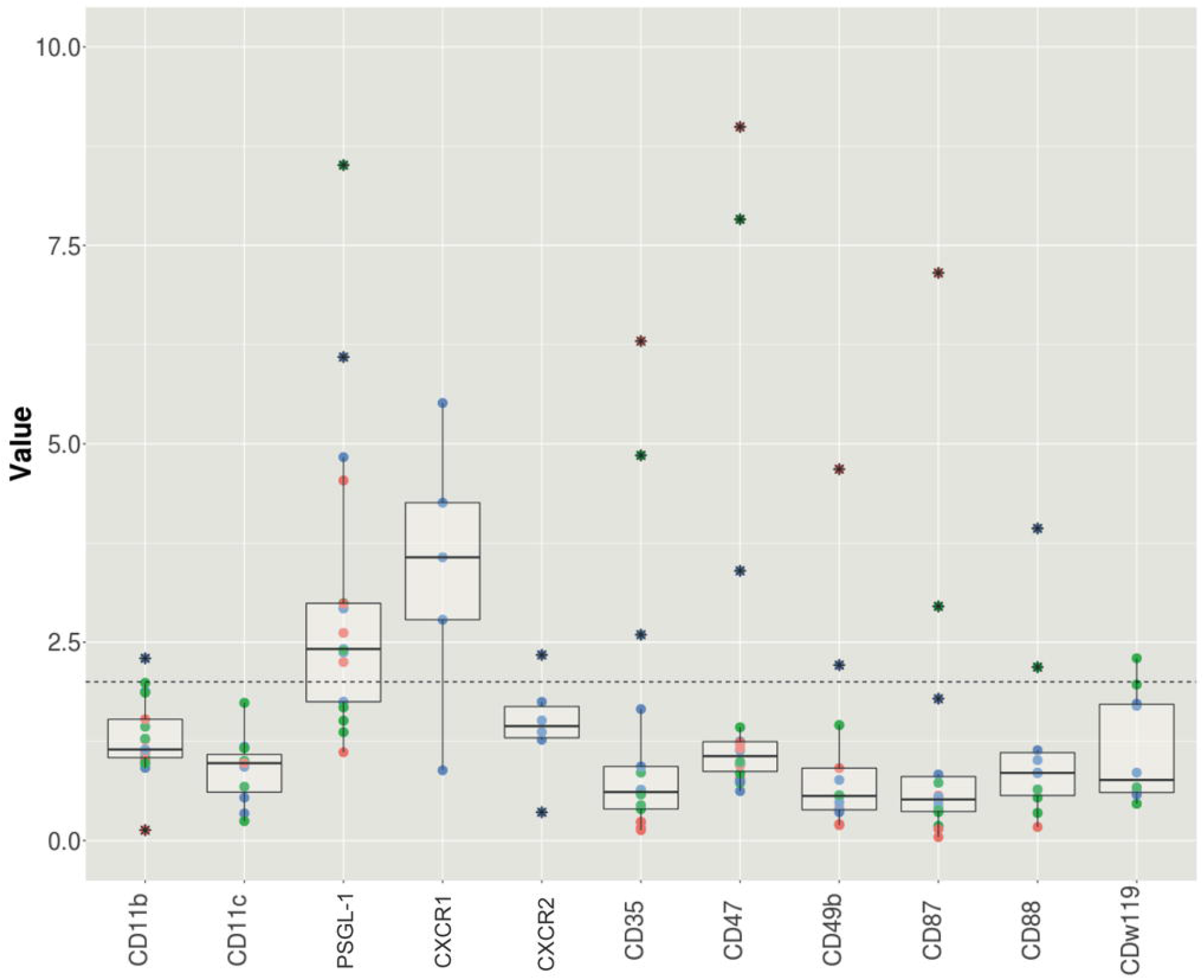
binding assay of the ≤ 3kDa fraction of the culture medium of strain AB1.13Δ*oahA*#76. Lymphocytes, monocytes, and neutrophils are represented with red, green, and blue dots, respectively, * represent outliers. Y-axis represent the inverted geometric mean of fluorescence, the X-axis represent the used receptors in the CBA, data points above the dotted line (2) are scored as positive for binding of molecules from the culture medium. Samples were tested in 3 independent experiments.

### Effect of PptA deletion on production of immune-competing components

We next investigated if molecules competing for binding with immune receptors produced by JN24.6 (Δ*laeA*) are synthesized via the NRPS or PKS pathway. Inactivation of *pptA* abolishes the production of secondary metabolites via the NRPS and PKS pathway. Strains lacking *pptA* require culture medium containing siderophores for growth, as deletion of *pptA* leads to impaired siderophore biosynthesis (45). In subsequent experiments siderophore medium was used to culture the Δ*pptA* strains as well as the other strains. Absence of *pptA* in N402 (JP1.1) did not affect medium acidification after 72 hours of growth in siderophore medium, while absence of medium acidification was observed for the Δ*laeA* strain (JN24.6) and the Δ*laeA*Δ*pptA* strain (MA870.1) (Table 3).

When grown in regular minimal medium, molecules competing for binding for the CD88, PSGL-1, CXCR1 and CXCR2 receptor are secreted into the culture medium by JN24.6 (Figure 4E). However, when this strain is grown in siderophore medium we only detected molecules competing for binding to the PSGL-1 receptor and no molecules competing for binding to the CD88, CXCR1 and CXCR2 receptor were produced (Figure 8B). Also, in strains lacking *pptA* (JP1.1) and both *pptA* and *laeA* (MA870.1) no competing molecules for CD88, CXCR1 and CXCR2 were detected (Figure 8A and C, respectively). Furthermore, the protein profile of the strains lacking *pptA* are rather different as compared to the control JN24.6. For example, strain JP1.1 produces proteins around 70 and 55 kDa that appear to be absent in the other strains (Figure 9). These results suggest that the use of siderophore medium inhibits the production of competing molecules for CD88, CXCR1 and CXCR2 receptor, while competing molecules for PSGL-1 were not influenced by the presence of siderophore medium (Figure 8).

**Figure 8.**
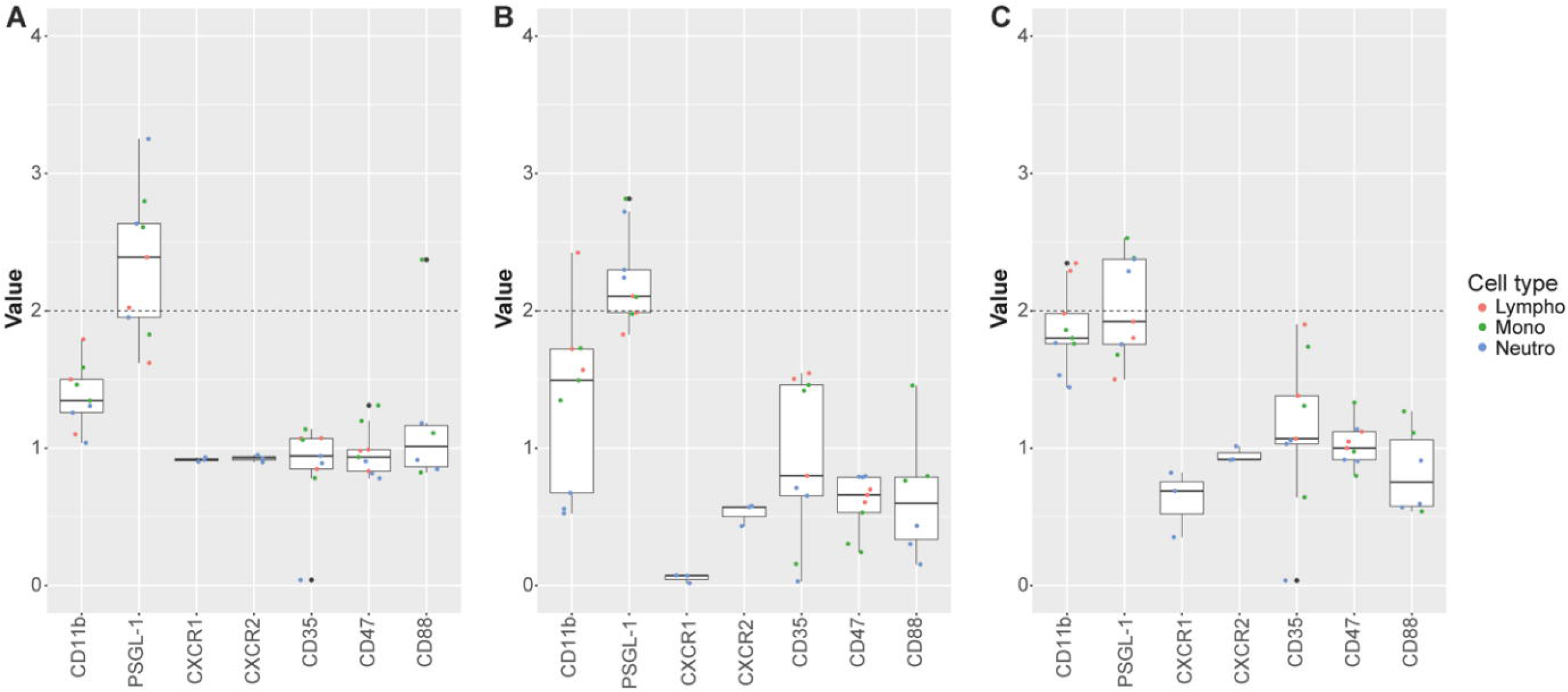
binding assay of the ≤ 3kDa fraction of the culture medium of strains JP1.1 (A), JN24.6 (B) and MA870.1 (C). Lymphocytes, monocytes, and neutrophils are represented with red, green, and blue dots, respectively, * represent outliers. Y-axis represent the inverted geometric mean of fluorescence, the X-axis represent the used receptors in the CBA, data points above the dotted line (2) are scored as positive for binding of molecules from the culture medium.

**Figure 9.**
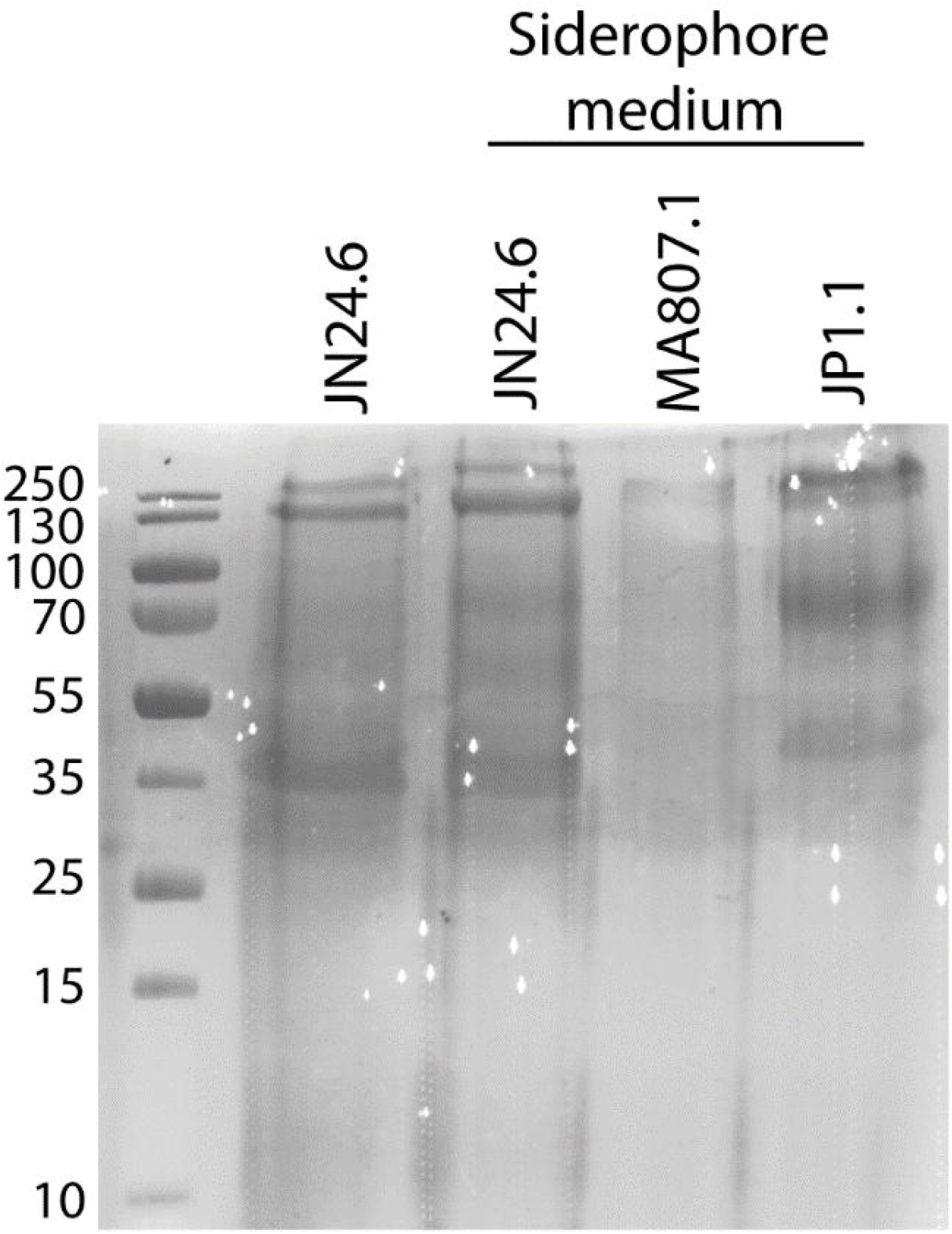
CBB stained SDS PAA gels of mixed maltose and xylose culture media after 72 h of growth of *A. niger* JN24.6 and of JN24.6, MA871.1 and JP1.1 after growth in siderophore medium.

## Discussion

Here we showed that culture media of *A. fumigatus* Af293 (46,47), *A. tubingensis* (48) and *A. niger* N402 and the non-acidifying *A. niger* strain D15#26 (49,50) contain ≤ 3 kDa compounds that could compete for binding with antibodies to human cellular receptors that have been related to immune recognition, activation, or modulation. Components within the culture media with competing activity were identified for the CD13, CD29, CD45, CD47, CD88, CD99, CD141, PSGL-1, CXCR1, CXCR2, CD191, CD192, and Siglec-9 receptors. The possible competing activity for PSGL-1, CXCR1, and CXCR2 was shared between the four strains. These receptors play a role in the recruitment of leukocytes (PSGL-1) and neutrophils (CXCR1 and CXCR2) to the site of infection. The immunological role of these and the other responsive receptors is well described (Supplementary Table 4) but to our knowledge none of them have been associated to fungal infections. The results of the CBA were in general quite variable, which is most likely due to the fact that for each experiment cells from different donors were used, as immune responses can vary per individual.

Initially, we hypothesized that fungal proteins were responsible for the competing activity with these receptors as has been described for S. *aureus* (10). Furthermore, a group of well-studied proteins in fungi were described as fungal immunomodulatory proteins (FIPs). Currently, more than 38 FIPs have been identified in different fungal species (51). FIPs are subdivided in 5 different groups, of which the Fve-type FIPs, small proteins around 13 kDa, and Cerato-type FIPs are most studied. The two groups can be identified by Pfam domain PF09259 and PF07249, respectively (51). We detected no Fve-type FIPs genes in the genome of *A. niger*, but we did detect a Cerato-type FIPs gene. A secreted serine protease (An02g01550) has a PF07249 domain. A more in-depth analysis of the genome of *A. niger* could indicate more possible FIPs. Nevertheless, no competing activity was recovered after protein purification using affinity chromatography, indicating that the competing activity is not due to fungal proteins. Filtration studies showed that molecules ≤ 3 kDa were involved. Such molecules can be peptides, carbohydrates, or secondary metabolites. The fact that we mainly detect small molecules suggests that Aspergilli might use also a different strategy for immune evasion. Further characterization and purification was performed using the *A. niger* D15#26 ≤ 3 kDa fractions. The molecules could not be heat inactivated. They bound to a C18 Sep-pak column and could be eluted with 10 - 50 % methanol, but were not extracted using ethyl acetate, suggesting that these molecules were relatively hydrophilic. The latter fractions contained a variety of low molecular weight molecules as shown by LC-MS.

Next to the characteristics of the small molecules produced, the mutations in the D15#26 responsible for the productions of the small molecules with competing activity for the immune receptors CD88, PSGL-1, CXCR1 and CXCR2 were assessed as well. Experimental data showed that strains with an inactive *pyrG* and / or *prtT* but with an intact *laeA* were not producing binding compounds except for PSGL-1. Preliminary CBA data with a Δ*prtT* strain (AF11#56, a derivative of AB4.1) underscored that deletion of PrtT did not lead to the production of CD88, PSGL-1, CXCR1, and CXCR2 binding compounds (Supplementary Figure 8). In contrast, gene *laeA* was shown to have a role as a repressor of production of competing compounds of CD88, CXCR1 and CXCR2. A Δ*laeA* strain did not acidify the culture medium like D15#26 and produced CD88, PSGL-1, CXCR1, and CXCR2 competing activity. On the other hand, the acidifying *laeA* complemented D15#26 strain only produced competing activity with PSGL-1, showing that the global regulator of secondary metabolism LaeA is a repressor of production of the competing activities to CD88, CXCR1, and CXCR2 (Figure 9). The finding that LaeA can act as a repressor was previously reported in *A. nidulans* and *A. niger*. These *laeA*-deficient strains showed increased secretion of an uncharacterized secondary metabolite (26) and aspernigrin (30), respectively. In addition, loss of *laeA* inhibits synthesis of sterigmatocystin and penicillin in *A. nidulans*, lovastatin in *A. terreus*, and gliotoxin in *A. fumigatus* (34,52).

Cultivation at neutral pH (between 5 and 6) is crucial for the production of molecules competing for binding to CXCR1 in *A. niger*. This was based on the fact that two non-acidifying strains, D15#26 and Δ*oahA* produced components that interacted with this receptor. Similar results were obtained in preliminary experiments with the non-acidifying *crzA* mutant of *A. niger* in an AB4.1 background (Supplementary Figure 9). CrzA is a transcription factor of the calcium / calcineurin pathway, involved in fungal morphogenesis, virulence, and ion tolerance (53). This shows that the competing activity for CXCR1 is only produced under neutral pH and its production depends indirectly on LaeA, as it is secreted under all non-acidifying conditions tested, while the mechanism underlying repression of CXCR2 and CD88 competing activity is linked to the presence of LaeA (Figure 10).

**Figure 10.**
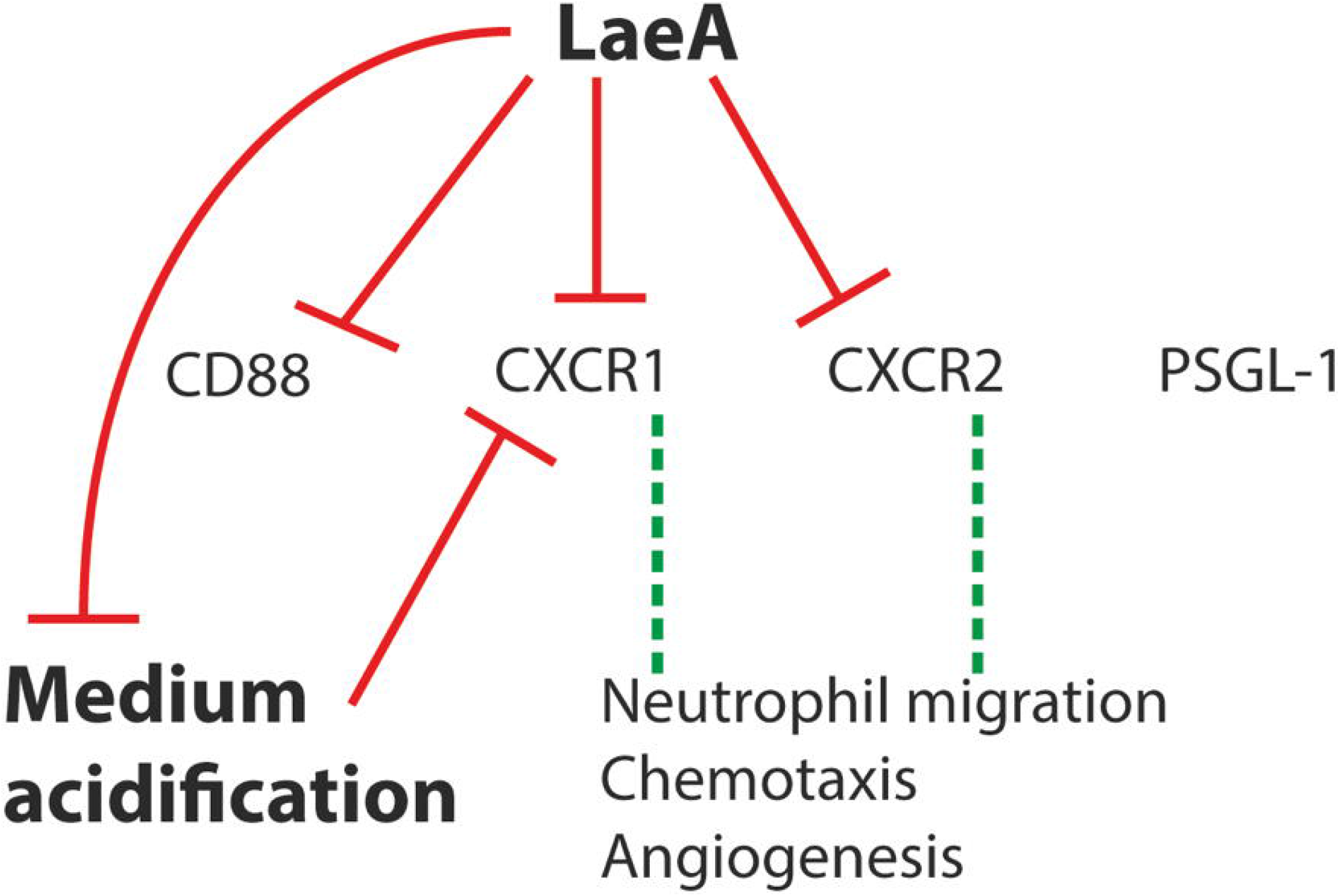
Model of the effectors and their role in the production of components competing with binding to the CD88, CXCR1, CXCR2 and PSGL-1 receptor.

Immune receptors CXCR1 and CXCR2 are better known as CXCR1 and CXCR2. They are chemokine receptors belonging to the G-protein-coupled receptor (GPCR) family. Ligands binding to CXCR1 and CXCR2 include IL-8, NAP-2, GCP-2, and GRO-α, β, γ (53–55). Activation of CXCR1 and CXCR2 mediates neutrophil migration and chemotaxis and favor angiogenesis (56). Both receptors are present on neutrophils, are closely related, and generally interact with similar ligands, but not necessarily with the same affinity (57–59). The observation that the CXCR1-interactive compound produced under non-acidifying conditions is not reacting with CXCR2 suggests the presence of two different compounds binding specifically to each receptor (Table 4).

**Table 4.** Summary of receptors with competition of binding from the fungal supernatant with antibodies

|  | A. niger |  |  |  |  |  |  |  |  | A. fumigatus |  | A. tubingensis |  |
| --- | --- | --- | --- | --- | --- | --- | --- | --- | --- | --- | --- | --- | --- |
|  | N402 |  | D15#26 |  | JN22 .7 | JN21 .1 | JN24 .6 | JN23 .1 | ΔoahA | Af293 |  |  |  |
|  | Whole fraction | ≤ 3k Da | Whole fraction | ≤ 3k Da | ≤ 3kDa | ≤ 3kDa | ≤ 3kDa | ≤ 3kDa | ≤ 3kDa | Whole fraction | ≤ 3k Da | Whole fraction | ≤ 3k Da |
| CD11b |  |  |  |  | X |  |  |  |  |  |  |  |  |
| CD47 |  |  | X |  | X |  |  |  |  |  |  |  | X |
| CD88 |  |  |  | X |  |  | X |  |  |  |  |  |  |
| CD141 |  |  | X |  |  |  |  |  |  |  |  |  |  |
| PSGL-1 | X | X | X | X | X | X | X | X | X | X | X | X | X |
| CXCR1 |  |  | X | X |  | X | X |  | X | X | X |  |  |
| CXCR2 |  |  | X | X |  | X | X |  |  | X | X |  |  |
| CD191 |  |  |  |  |  |  |  |  |  | X | X |  |  |
| CD192 |  |  |  |  |  |  |  |  |  |  |  |  | X |

Together, our results show that we are dealing with small hydrophilic molecules which could be secondary metabolites, carbohydrates, and / or small peptides that could be responsible for competition for binding of antibodies to cellular receptors. *Aspergilli* secrete polyketides (PKS), non-ribosomal peptides (NRPS), terpenes, and indole alkaloids as main groups of secondary metabolites (40). Deletion of the *pptA* gene abolishes secondary metabolite production via the PKS and NRPS pathway and to be able to grow these strains need siderophore medium (37). Growth of the Δ*laeA* strain in siderophore medium did not alter medium acidification (Table 3), but the molecules competing for binding with the CD88, CXCR1 and CXCR2 were not produced (Figure 8B). This indicates that the addition of medium containing siderophore produced by *A. niger* affects the production of these molecules suggesting that iron limitation might be involved in their regulation. In line with these results, we also do not see the production of these molecules in the Δ*pptA* strain and Δ*laeA*Δ*pptA* strain (Figure 8A and C, respectively). We therefore could not determine if the small molecules were indeed produced via a PKS and/or NRPS pathways and which could confirm they are secondary metabolites. Future research is needed to elucidate if the molecules competing for binding are indeed secondary metabolites and whether they are produced via a PKS, NRPS or other pathways involved in synthesis of secondary metabolites.

Genes required for the biosynthesis of these metabolites are usually located in gene clusters, with some exceptions like in *A. nidulans* where two gene clusters located on separate chromosomes are required for the production of meroterpenoids austinol and dehydroaustinol (60). *A. niger* D15#26 produces more responsive molecules when compared to its wild type strain N402. D15#26 contains a variety of mutations including a mutation in *laeA* that controls production of secondary metabolites like sterigmatocystin, penicillin, and lovastatin (26,30). The competing components produced by D15#26 are expressed in a LaeA-dependent manner. *A. niger* ATCC 1015 is predicted to have 81 gene clusters associated with secondary metabolite production (61) and 145 secondary metabolites have been identified from the *Aspergillus Nigri* section. Many of these compounds (e.g. ochratoxin A, naptho-γ-pyrones, bocoumarins) are found in both *A. niger and A. tubingensis* (23). As described by (61), *A. fumigatus* Af293 contains 39 secondary metabolite gene clusters, while at least 226 *A. fumigatus* secondary metabolites have been reported, some of them associated with virulence (62,63). The fact that components produced by D15#26 (lacking LaeA) and *A. fumigatus* and *A. tubingensis* (both containing LaeA) compete for to the same set of receptors (PSGL-1, CXCR1, and CXCR2) indicates that we are dealing with a variety of molecules that are regulated differently but might be produced by orthologous gene clusters. By using Anti-SMASH prediction software, we detected 88 gene clusters in the genome of *A. tubingensis* CBS134.48. Of these, 30 gene clusters showed ≥ 75 % homology at the amino acid level when compared to *A. niger* ATCC 1015. 4 gene clusters showed 50 % homology at the amino acid level when compared to *A. fumigatus* Af293. The 4 gene clusters having similarity between *A. tubingensis* CBS134.48 and *A. fumigatus* Af293 were also found in *A. niger* ATCC 1015. In this case a similarity ≥ 57 % at the amino acid level was found in the latter strains. Shared clusters were assigned and predicted in *A. tubingensis* as Cluster 4 (non-ribosomal peptide), Cluster 18 (type I polyketide synthase), Cluster 27 (terpene), and Cluster 79 (other). Cluster 4 showed similarity with *A. fumigatus* Afu1g10380 (nrps1 / pes1), while Cluster 18 had similarity with *A. fumigatus* Afu2g01290, Cluster 27 with No PKS or NRPS backbone 6 and cluster 79 with No PKS or NRPS backbone 2. These clusters are of interest for further analysis with respect to immune receptor binding.

Identification of the small compounds secreted by *A. niger laeA* mutant strains might result in novel therapeutic agents. Furthermore, absence of medium acidification might also explain the secretion of immune-modulatory components of the pathogen *A. fumigatus*. In this study it was shown that this species produced components ≤ 3 kDa that might compete with antibodies for interaction with receptors PSGL-1, CXCR1, CXCR2, CD192, CD99, CD45, CD47, and CD29. Even though several purification methods and a preparative HPLC was done, we were unable to identify the molecules which compete for binding with the immune receptors. More research should be done to identify these molecules and determine their role in infection.

Interestingly, *A. fumigatus* increases the pH of the culture medium to 8. This increase may be responsible for the production of the binding compounds. Notably, (20,64) reported that secretion of serine protease (Alp1), metalloproteases (Mep1), and leucine aminopeptidases (Lap1 and Lap2) by *A. fumigatus* were favoured at pH between 7 −7.5 but undetected at pH 3.5 (20,64). Secretion of Alp1 is related with immune evasion as it degrades human complements proteins C3, C4, and C5 (12). Possibly, PacC that is required for alkaline adaption and implicated as another global regulator of secondary metabolite production (34,65) plays an important role in the production of these compounds.

## Declarations

### Availability of data and material

All data and material belonging to this study are available on request

### Code availability

software application used in this study is freely available and sources are described

### Authors’ contributions

NE, EK, JvN, MA performed experiments and provided strains. PJH, JvS, AR, PP, HW and HdC designed experiments. NE and EK wrote the original version of the manuscript. All authors edited the manuscript and approved the final version.

### Ethics approval

#### Consent to participate and publication

Written informed consent was obtained from all subjects according to the Declaration of Helsinki. Approval was obtained from the medical ethics committee of the University Medical Center Utrecht (Utrecht, The Netherlands).

## Acknowledgements

We thank Steven Braem, Annelies Smout and Tiemen Knoop for their help in initial experiments and Jelmer Hoeksma for his help with HPLC and LC-MS analysis. We thank Dr. Kok van Kessel for his help with the CBA-assay and the FACS analysis.

## References

(1) Levitz SM, Farrell TP. Human neutrophil degranulation stimulated by Aspergillus fumigatus. J Leukoc Biol 1990 Feb;47(2):170–175.

(2) Fietta A, Sacchi F, Mangiarotti P, Manara G, Gialdroni Grassi G. Defective phagocyte Aspergillus killing associated with recurrent pulmonary Aspergillus infections. Infection 1984 Jan-Feb;12(1):10–13.

(3) Kan VL, Bennett JE. Beta 1,4-oligoglucosides inhibit the binding of Aspergillus fumigatus conidia to human monocytes. J Infect Dis 1991 May;163(5):1154–1156.

(4) Li SS, Kyei SK, Timm-McCann M, Ogbomo H, Jones GJ, Shi M, et al. The NK receptor NKp30 mediates direct fungal recognition and killing and is diminished in NK cells from HIV-infected patients. Cell Host Microbe 2013 Oct 16,;14(4):387–397.

(5) Shoham S, Levitz SM. The immune response to fungal infections. Br J Haematol 2005 Jun;129(5):569–582.

(6) Mircescu MM, Lipuma L, van Rooijen N, Pamer EG, Hohl TM. Essential role for neutrophils but not alveolar macrophages at early time points following Aspergillus fumigatus infection. J Infect Dis 2009 Aug 15,;200(4):647–656.

(7) Werner JL, Metz AE, Horn D, Schoeb TR, Hewitt MM, Schwiebert LM, et al. Requisite role for the dectin-1 beta-glucan receptor in pulmonary defense against Aspergillus fumigatus. J Immunol 2009 Apr 15,;182(8):4938–4946.

(8) Brown GD. Dectin-1: a signalling non-TLR pattern-recognition receptor. Nat Rev Immunol 2006 Jan;6(1):33–43.

(9) Swidergall M, Solis NV, Lionakis MS, Filler SG. EphA2 is an epithelial cell pattern recognition receptor for fungal β-glucans. Nature Microbiology 2018 -01;3(1):53–61.

(10) Bestebroer J, Poppelier, Miriam J. J. G., Ulfman LH, Lenting PJ, Denis CV, van Kessel, Kok P. M., et al. Staphylococcal superantigen-like 5 binds PSGL-1 and inhibits P-selectin-mediated neutrophil rolling. Blood 2007 Apr 01,;109(7):2936–2943.

(11) Shende R, Wong SSW, Rapole S, Beau R, Ibrahim-Granet O, Monod M, et al. Aspergillus fumigatus conidial metalloprotease Mep1p cleaves host complement proteins. J Biol Chem 2018 10 05,;293(40):15538–15555.

(12) Behnsen J, Lessing F, Schindler S, Wartenberg D, Jacobsen ID, Thoen M, et al. Secreted Aspergillus fumigatus protease Alp1 degrades human complement proteins C3, C4, and C5. Infect Immun 2010 Aug;78(8):3585–3594.

(13) Tsang A, Butler G, Powlowski J, Panisko EA, Baker SE. Analytical and computational approaches to define the Aspergillus niger secretome. Fungal Genet Biol 2009 Mar;46 Suppl 1:S153–S160.

(14) Punt PJ, van Biezen N, Conesa A, Albers A, Mangnus J, van den Hondel C. Filamentous fungi as cell factories for heterologous protein production. Trends Biotechnol 2002 May;20(5):200–206.

(15) Medina ML, Haynes PA, Breci L, Francisco WA. Analysis of secreted proteins from Aspergillus flavus. Proteomics 2005 Aug;5(12):3153–3161.

(16) Machida M, Asai K, Sano M, Tanaka T, Kumagai T, Terai G, et al. Genome sequencing and analysis of Aspergillus oryzae. Nature 2005 Dec 22,;438(7071):1157–1161.

(17) Brakhage AA, Schroeckh V. Fungal secondary metabolites - strategies to activate silent gene clusters. Fungal Genet Biol 2011 Jan;48(1):15–22.

(18) Lu X, Sun J, Nimtz M, Wissing J, Zeng A, Rinas U. The intra- and extracellular proteome of Aspergillus niger growing on defined medium with xylose or maltose as carbon substrate. Microb Cell Fact 2010 Apr 20,;9:23.

(19) Giorni P, Battilani P, Pietri A, Magan N. Effect of aw and CO2 level on Aspergillus flavus growth and aflatoxin production in high moisture maize post-harvest. Int J Food Microbiol 2008 Feb 29,;122(1-2):109–113.

(20) Sriranganadane D, Waridel P, Salamin K, Reichard U, Grouzmann E, Neuhaus J, et al. Aspergillus protein degradation pathways with different secreted protease sets at neutral and acidic pH. J Proteome Res 2010 Jul 02,;9(7):3511–3519.

(21) Wongwicharn A, McNeil B, Harvey LM. Effect of oxygen enrichment on morphology, growth, and heterologous protein production in chemostat cultures of Aspergillus niger B1-D. Biotechnol Bioeng 1999 Nov 20,;65(4):416–424.

(22) Braaksma M, Martens-Uzunova ES, Punt PJ, Schaap PJ. An inventory of the Aspergillus niger secretome by combining in silico predictions with shotgun proteomics data. BMC Genomics 2010 Oct 19,;11:584.

(23) Nielsen KF, Mogensen JM, Johansen M, Larsen TO, Frisvad JC. Review of secondary metabolites and mycotoxins from the Aspergillus niger group. Anal Bioanal Chem 2009 Nov;395(5):1225–1242.

(24) Budak SO, Zhou M, Brouwer C, Wiebenga A, Benoit I, Di Falco M, et al. A genomic survey of proteases in Aspergilli. BMC Genomics 2014 Jun 25,;15:523.

(25) Sharon H, Amar D, Levdansky E, Mircus G, Shadkchan Y, Shamir R, et al. PrtT-regulated proteins secreted by Aspergillus fumigatus activate MAPK signaling in exposed A549 lung cells leading to necrotic cell death. PLoS ONE 2011 Mar 11,;6(3):e17509.

(26) Bok JW, Keller NP. LaeA, a regulator of secondary metabolism in Aspergillus spp. Eukaryotic Cell 2004 Apr;3(2):527–535.

(27) Punt PJ, Schuren FHJ, Lehmbeck J, Christensen T, Hjort C, van den Hondel, Cees A. M. J. J. Characterization of the Aspergillus niger prtT, a unique regulator of extracellular protease encoding genes. Fungal Genet Biol 2008 Dec;45(12):1591–1599.

(28) Hagag S, Kubitschek-Barreira P, Neves GWP, Amar D, Nierman W, Shalit I, et al. Transcriptional and proteomic analysis of the Aspergillus fumigatus ΔprtT protease-deficient mutant. PLoS ONE 2012;7(4):e33604.

(29) Sharon H, Hagag S, Osherov N. Transcription factor PrtT controls expression of multiple secreted proteases in the human pathogenic mold Aspergillus fumigatus. Infect Immun 2009 Sep;77(9):4051–4060.

(30) Niu J, Arentshorst M, Nair PDS, Dai Z, Baker SE, Frisvad JC, et al. Identification of a Classical Mutant in the Industrial Host Aspergillus niger by Systems Genetics: LaeA Is Required for Citric Acid Production and Regulates the Formation of Some Secondary Metabolites. G3 (Bethesda) 2015 Nov 13,;6(1):193–204.

(31) Andersen MR, Lehmann L, Nielsen J. Systemic analysis of the response of Aspergillus niger to ambient pH. Genome Biol 2009;10(5):R47.

(32) Karaffa L, Kubicek CP. Aspergillus niger citric acid accumulation: do we understand this well working black box? Appl Microbiol Biotechnol 2003 May;61(3):189–196.

(33) Ruijter GJG, van de Vondervoort, Peter J. I., Visser J. Oxalic acid production by Aspergillus niger: an oxalate-non-producing mutant produces citric acid at pH 5 and in the presence of manganese. Microbiology (Reading, Engl) 1999 Sep;145 (Pt 9):2569–2576.

(34) Brakhage AA. Regulation of fungal secondary metabolism. Nat Rev Microbiol 2013 Jan;11(1):21–32.

(35) Keszenman-Pereyra D, Lawrence S, Twfieg M, Price J, Turner G. The npgA/ cfwA gene encodes a putative 4’-phosphopantetheinyl transferase which is essential for penicillin biosynthesis in Aspergillus nidulans. Curr Genet 2003 Jun;43(3):186–190.

(36) Jørgensen TR, Park J, Arentshorst M, van Welzen AM, Lamers G, Vankuyk PA, et al. The molecular and genetic basis of conidial pigmentation in Aspergillus niger. Fungal Genet Biol 2011 May;48(5):544–553.

(37) Márquez-Fernández O, Trigos A, Ramos-Balderas JL, Viniegra-González G, Deising HB, Aguirre J. Phosphopantetheinyl transferase CfwA/NpgA is required for Aspergillus nidulans secondary metabolism and asexual development. Eukaryotic Cell 2007 Apr;6(4):710–720.

(38) Johns A, Scharf DH, Gsaller F, Schmidt H, Heinekamp T, Straßburger M, et al. A Nonredundant Phosphopantetheinyl Transferase, PptA, Is a Novel Antifungal Target That Directs Secondary Metabolite, Siderophore, and Lysine Biosynthesis in Aspergillus fumigatus and Is Critical for Pathogenicity. mBio 2017 07 18,;8(4).

(39) Dewick PM. Medicinal natural products: a biosynthetic approach. 3rd ed. Chichester: John Wiley & Sons; 2009.

(40) Keller NP, Turner G, Bennett JW. Fungal secondary metabolism - from biochemistry to genomics. Nat Rev Microbiol 2005 Dec;3(12):937–947.

(41) Arentshorst M, Niu J, Ram A. Efficient Generation of Aspergillus niger Knock Out Strains by Combining NHEJ Mutants and a Split Marker Approach.; 2015. p. 263–272.

(42) Bos CJ, Debets AJ, Swart K, Huybers A, Kobus G, Slakhorst SM. Genetic analysis and the construction of master strains for assignment of genes to six linkage groups in Aspergillus niger. Curr Genet 1988 Nov;14(5):437–443.

(43) Punt PJ, Oliver RP, Dingemanse MA, Pouwels PH, van den Hondel, C. A. Transformation of Aspergillus based on the hygromycin B resistance marker from Escherichia coli. Gene 1987;56(1):117–124.

(44) de Vries RP, Riley R, Wiebenga A, Aguilar-Osorio G, Amillis S, Uchima CA, et al. Comparative genomics reveals high biological diversity and specific adaptations in the industrially and medically important fungal genus Aspergillus. Genome Biol 2017 02 14,;18(1):28.

(45) Oberegger H, Eisendle M, Schrettl M, Graessle S, Haas H. 4’-phosphopantetheinyl transferase-encoding npgA is essential for siderophore biosynthesis in Aspergillus nidulans. Curr Genet 2003 Dec;44(4):211–215.

(46) Pain A, Woodward J, Quail MA, Anderson MJ, Clark R, Collins M, et al. Insight into the genome of Aspergillus fumigatus: analysis of a 922 kb region encompassing the nitrate assimilation gene cluster. Fungal Genet Biol 2004 Apr;41(4):443–453.

(47) Nierman WC, Pain A, Anderson MJ, Wortman JR, Kim HS, Arroyo J, et al. Genomic sequence of the pathogenic and allergenic filamentous fungus Aspergillus fumigatus. Nature 2005 Dec 22,;438(7071):1151–1156.

(48) Bathoorn E, Escobar Salazar N, Sepehrkhouy S, Meijer M, de Cock H, Haas P. Involvement of the opportunistic pathogen Aspergillus tubingensis in osteomyelitis of the maxillary bone: a case report. BMC Infect Dis 2013 Feb 01,;13:59.

(49) Gordon CL, Khalaj V, Ram AFJ, Archer DB, Brookman JL, Trinci APJ, et al. Glucoamylase::green fluorescent protein fusions to monitor protein secretion in Aspergillus niger. Microbiology (Reading, Engl) 2000 Feb;146 (Pt 2):415–426.

(50) Pham TA, Berrin JG, Record E, To KA, Sigoillot J. Hydrolysis of softwood by Aspergillus mannanase: role of a carbohydrate-binding module. J Biotechnol 2010 Aug 02,;148(4):163–170.

(51) Liu Y, Bastiaan-Net S, Wichers HJ. Current Understanding of the Structure and Function of Fungal Immunomodulatory Proteins. Front Nutr 2020 -8-18;7.

(52) Bok JW, Noordermeer D, Kale SP, Keller NP. Secondary metabolic gene cluster silencing in Aspergillus nidulans. Mol Microbiol 2006 Sep;61(6):1636–1645.

(53) de Castro PA, Chen C, de Almeida, Ricardo Sérgio Couto, Freitas FZ, Bertolini MC, Morais ER, et al. ChIP-seq reveals a role for CrzA in the Aspergillus fumigatus high-osmolarity glycerol response (HOG) signalling pathway. Mol Microbiol 2014 Nov;94(3):655–674.

(54) Lee J, Horuk R, Rice GC, Bennett GL, Camerato T, Wood WI. Characterization of two high affinity human interleukin-8 receptors. J Biol Chem 1992 Aug 15,;267(23):16283–16287.

(55) Ahuja SK, Murphy PM. The CXC chemokines growth-regulated oncogene (GRO) alpha, GRObeta, GROgamma, neutrophil-activating peptide-2, and epithelial cell-derived neutrophil-activating peptide-78 are potent agonists for the type B, but not the type A, human interleukin-8 receptor. J Biol Chem 1996 Aug 23,;271(34):20545–20550.

(56) Stillie R, Farooq SM, Gordon JR, Stadnyk AW. The functional significance behind expressing two IL-8 receptor types on PMN. J Leukoc Biol 2009 Sep;86(3):529–543.

(57) Cummings CJ, Martin TR, Frevert CW, Quan JM, Wong VA, Mongovin SM, et al. Expression and function of the chemokine receptors CXCR1 and CXCR2 in sepsis. J Immunol 1999 Feb 15,;162(4):2341–2346.

(58) Cerretti DP, Kozlosky CJ, Vanden Bos T, Nelson N, Gearing DP, Beckmann MP. Molecular characterization of receptors for human interleukin-8, GRO/melanoma growth-stimulatory activity and neutrophil activating peptide-2. Mol Immunol 1993 Mar;30(4):359–367.

(59) Wolf M, Delgado MB, Jones SA, Dewald B, Clark-Lewis I, Baggiolini M. Granulocyte chemotactic protein 2 acts via both IL-8 receptors, CXCR1 and CXCR2. Eur J Immunol 1998 Jan;28(1):164–170.

(60) Lo H, Entwistle R, Guo C, Ahuja M, Szewczyk E, Hung J, et al. Two separate gene clusters encode the biosynthetic pathway for the meroterpenoids austinol and dehydroaustinol in Aspergillus nidulans. J Am Chem Soc 2012 Mar 14,;134(10):4709–4720.

(61) Inglis DO, Binkley J, Skrzypek MS, Arnaud MB, Cerqueira GC, Shah P, et al. Comprehensive annotation of secondary metabolite biosynthetic genes and gene clusters of Aspergillus nidulans, A. fumigatus, A. niger and A. oryzae. BMC Microbiol 2013 Apr 26,;13:91.

(62) Frisvad JC, Rank C, Nielsen KF, Larsen TO. Metabolomics of Aspergillus fumigatus. Med Mycol 2009;47 Suppl 1:53.

(63) Latgé JP. Aspergillus fumigatus and aspergillosis. Clin Microbiol Rev 1999 Apr;12(2):310–350.

(64) Kniemeyer O. Proteomics of eukaryotic microorganisms: The medically and biotechnologically important fungal genus Aspergillus. Proteomics 2011 Aug;11(15):3232–3243.

(65) Tilburn J, Sarkar S, Widdick DA, Espeso EA, Orejas M, Mungroo J, et al. The Aspergillus PacC zinc finger transcription factor mediates regulation of both acid- and alkaline-expressed genes by ambient pH. EMBO J 1995 Feb 15,;14(4):779–790.

(66) Mattern IE, van Noort JM, van den Berg P, Archer DB, Roberts IN, van den Hondel, C. A. Isolation and characterization of mutants of Aspergillus niger deficient in extracellular proteases. Mol Gen Genet 1992 Aug;234(2):332–336.

(67) Li A, Pfelzer N, Zuijderwijk R, Punt P. Enhanced itaconic acid production in Aspergillus niger using genetic modification and medium optimization. BMC Biotechnol 2012 Aug 27,;12:57.

(68) A. W. C. Franken, B. C. Lokman, van den Hondel, C. A. M. J. J., S. de Weert. Peroxidase production in A. niger for white biotechnologyLeiden University; 2017.

