## Supplementary information for "LaeA-dependent production of small molecules of *Aspergillus niger* that compete with specific antibodies that bind to human immune receptors"

**Supplementary Table 1.** Overview of the primers used in this study

| **Oligo** | **Sequence 5'-3'** | **Used for** |
|---|---|---|
| **Deletion of pptA::hygR with split marker method** | | |
| underlined: overlapping sequence used in fusion PCR | | |
| pptAP1f | GATCCTGGCCACCCTTCTTT | pptA 5' flank + pptA-hygR 5' SM fragment |
| pptAP2r | CAATTCCAGCAGCGGCTTTGTGGGTTGGTTGGGTTGGGTA | pptA 5' flank |
| pptAP3f | ACACGGCACAATTATCCATCGGATATCTATTTATTTAACTATGAGTG | pptA 3' flank |
| pptAP4r | AACGGCCGTCAAGGCTTT | pptA 3' flank + pptA-hygR 3' SM fragment |
| hygP6f | AAGCCGCTGCTGGAATTGGGCTCTGAGGTGCAGTGGAT | hygR 5' fragment |
| hygP9r | GGCGTCGGTTTCCACTATC | hygR 5' fragment + pptA-hygR 5' SM fragment |
| hygP7r | CGATGGATAATTGTGCCGTGTTGGGTGTTACGGAGCATTCA | hygR 3' fragment |
| hygP8f | AAAGTTCGACAGCGTCTCC | hygR 3' fragment + pptA-hygR 3' SM fragment |
| hygP2f | CATGCATGGTTGCCTAGTGAA | Diagnostic PCR to confirm deletion |
| hygP5r | ATCCACTGCACCTCAGAGCC | Diagnostic PCR to confirm deletion |
| pptAP5f | AGGAAGGGCAAGTCGAGAGG | Diagnostic PCR to confirm deletion |
| pptAP6r | TTGAGGAGGTTGGAGGCTAG G | Diagnostic PCR to confirm deletion |
| pptAP7f | TGGGAGGGATTGAGGAGGTT | Diagnostic PCR to confirm deletion |
| pptAP8r | TCTTCCCTCCGTACTGCAGC | Diagnostic PCR to confirm deletion |

**Supplementary Table 2.** Volume per well used in the CBA. * Pre-dilutions (CD15 1:10, fMLP 1:100, CD32 1:10, CD44 1:5, CD16 1:40, CD45 1:10). The cell types where receptors are expressed are indicated by neutrophils (N), monocytes (M), and lymphocytes (L).

| Well | FITC | uL | Cell type | Clone/brand | PE | uL | Cell type | Clone/brand | APC | uL | Cell type | Clone/brand |
|---|---|---|---|---|---|---|---|---|---|---|---|---|
| 1 | CD11a | 0,5 | N, M,L | HI111, BD | CD32* | 1 | N, M,L | 7.3 /Fitzgerald | CD 10 | 1,5 | N, M,L | MEM-78, Invitrogen |
| 2 | CD15* | 1 | N | MMA, BD | CD35 | 2,5 | N, M,L | E11/ BD | CD 11c | 4 | N, M | B-ly6, BD |
| 3 | CD147 | 1,5 | N, M,L | HIM6, BD | CD44* | 1 | N, M,L | 515, BD | CD 11b | 4 | N, M,L | ICRF44, BD |
| 4 | CD18 | 1 | N, M,L | 6.7, BD | CD47 | 3 | N, M,L | B6H12, BD | CD 13 | 4 | N, M | WM15, BD |
| 5 | CD31 | 1,5 | N, M,L | WM59, BD | CD54 | 2 | N, M,L | HA58, BD | CD 14 | 4 | N, M | M5E2, BD |
| 6 | CD46 | 1 | N, M,L | E4.3, BD | CD58 | 1,5 | N, M,L | 1C3, BD | CD 29 | 1,5 | N, M,L | MAR4, BD |
| 7 | CD99 | 3 | M | Tü12,BD | CD321 | 0,75 | N, M,L | M.Ab.F11, BD | CD 16* | 1,5 | N, M,L | 3G8, BD |
| 8 | CD102 | 2 | M | aa191-218, BioCon | CD87 | 5 | N, M,L | VIM5, BD | CD 45* | 2 | N, M,L | HI30, BD |
| 9 | CD61 | 1,5 | M | 10.1, DB | CD89 | 2 | N, M | Mip8a, Abd-Serotec | CD 55 | 1 | N, M,L | IA10, BD |
| 10 | CD62L | 1,5 | N, M,L | Dreg-56, BD | CD63 | 1 | N, M | CLB-gran/12, Immuno Tech | CD 50 | 0,3 | N, M,L | CBR-IC3/1, BD |
| 11 | CD64 | 1,5 | M | VI-PL2, BD | CD114 | 1,5 | N, M | LMM741, BD | JAM-C | 2,5 | M,L | 208212, R&D |
| 12 | CD66b | 1,5 | N | G10F5, BD | CDw 119 | 1 | N, M | GIR-208, BD | Siglec-9 | 0,75 | N, M | 191240, R&D |
| 13 | CD43 | 1 | N, M,L | 6D269, Santa Cruz | CD132 | 2 | M | TUGh4, BD | CD 141 | 3 | M | 1A4, BD |
| 14 | CD66 | 1 | N, M | B1.1/CD66, BD | CD151 | 1,5 | M,L | 14A2.H1,BD | | | | |
| 15 | CD184 | 4 | M | 20102, R&D | PSGL-1 | 2 | N, M,L | KPL-1, BD | | | | |
| 16 | fMLP* | 1,5 | N, M | Rea169, MilBio | CD191 | 4 | M | 53504, R&D | | | | |

| | | | | | | | | |
|---|---|---|---|---|---|---|---|---|
| 17 | HLA-DR | 2 | M | G46-6,BD | CD192 | 2 | M | 48607, R&D |
| 18 | LTB4R | 2 | N, M | 202/7B1, Abd-Serotec | CXCR1 | 1, 5 | N | 42705.111, R&D |
| 19 | CD284 | 4 | M | HTA125, AbDSerotec | CXCR2 | 2, 5 | N | 48311.211, BD |
| 20 | CD120a | 4 | N, M | 16803.161, R&D | CD88 | 0, 3 | N, M | S5/1, Biolegend |
| 21 | CD120b | 3 | N, M,L | 22235.311, R&D | CD282 | 0, 5 | N, M | T2.5, EBioscience |
| 22 | CD9 | 2 | N, M,L | M-L13, BD | CD49b | 1 | N, M,L | 12F1, BD |
| 23 | TLR6 | 8 | N, M | 86B1153.2, Invitrogen | Dectin-1 | 4 | N, M | 15E2, BD |

BD (Becton, Dickinson and company, France), Santa Cruz (Dallas, USA), Abd-Serotec (MorphoSys ABD GmbH ABD Serotec, Düsseldorf, Germany), R&D (R&D Systems Inc, Minneapolis, USA), Fitzgerald (Fitzgerald Industries International, Acton, USA), ImmunoTech (Vaudreuil-Dorion, Canada), Biolegend (San Diego, USA), EBioscience (San Diego, USA), Invitrogen (Carlsbad, USA), MilBio (Miltenyi Biotec, Bergisch Gladbach, Germany), BioCon (BioConnect Life Sicence, Huissen, The Netherlands).

**Supplementary Table 3.** LC-MS and preparative HPLC gradient elution profile. Buffer A: 95 % MiliQ water, 5 % acetronitrile, and 0.1 % trifluoroacetic acid. Buffer B: 5 % MiliQ water, 95 % acetronitrile and 0.1 % trifluoroacetic acid.

| **Time (min)** | **Percentage Buffer B** |
|---|---|
| 0-5 | 0 |
| 5-80 | 0-60 |
| 80-85 | 60-100 |
| 85-90 | 100 |
| 90-95 | 100-0 |
| 95-100 | 0 |

**Supplementary Table 4.** Function and description of cellular receptors to which molecules from *Aspergillus* spp. supernatants bound.

| Receptor | Synonyms | Encodes | Function |
|---|---|---|---|
| PSGL-1 | P-selectin ligand | Glycoprotein-receptor for the cell adhesion molecules P- E- and L-selectins. | Leukocyte rolling and trafficking during inflammation. Activates platelets. |
| CXCR1 | CXCR1 | seven-transmembrane G-protein (67-70 kD) | Receptor to interleukin-8. Activation and chemotaxis of neutrophils. |
| CXCR2 | CXCR2 | seven-transmembrane G-protein (67-70 kD) | Receptor to interleukin-8. Activation and chemotaxis of neutrophils. |
| CD88 | C5aR | Single chain protein with seven membrane-spanning regions (43 kD). | Binds to complement component 5; stimulating chemotaxis, granule enzyme release, intracellular calcium release and superoxide anion production. |
| Siglec-9 | Sialic acid-binding Ig-like lectin-9 | Member of the Siglec subgroup of the immunoglobulin superfamily. | Negative regulation of T cell activation. Affects neutrophil apoptosis |
| CD99 | MIC2 | Single chain transmembrane protein (32 kD) | Involve in T-cell adhesion and spontaneous rosette formation with erythrocytes. |
| CD47 | Rh-associated protein | Integrin-associated protein (IAP). Member of the immunoglobulin superfamily (42-52 kD). | Cell addition and modulation of integrins. Associates with SIRP1a; involved in migration and phagocytosis. |
| CD45 | Leukocyte Common Antigen (LCA) | Single chain type I membrane glycoprotein, tyrosine-protein phosphatase C (180 – 240 KD) | Regulator of T-cell coactivation upon binding to DPP4. Also regulates cell growth, differentiation, cell cycle, and oncogenic transformation. |
| CD13 | Aminopeptidase N | Type II transmembrane glycoprotein, aminopeptidase N. | Antigen processing and cleavage of chemokines. |
| CD191 | CCR1 | C-C chemokine receptor type 1 (41 kD). | Inflammation and susceptibility to viruses and parasites. |
| CD192 | CKR2, CCR2A | Transmembrane glycoprotein and member of the G- | Role in inflammation. Induce infiltration of |

| | | | |
|---|---|---|---|
| | | protein coupled receptor 1 family. | macrophages. Associated with obesity. |
| CD141 | Thrombomodulin, TM | Single chain, type I membrane glycoprotein, also known as thrombomodulin. (75 kD) | Activation of protein C and reduction of thrombin. Anticoagulation function and embryonic and atherosclerotic plaque development. |
| CD29 | Integrin $\beta_1$ chain, ITGB1 | Type I glycoprotein (130 kD) | Acts as a fibronectin receptor and is involved in cell-cell and cell-matrix interactions. |

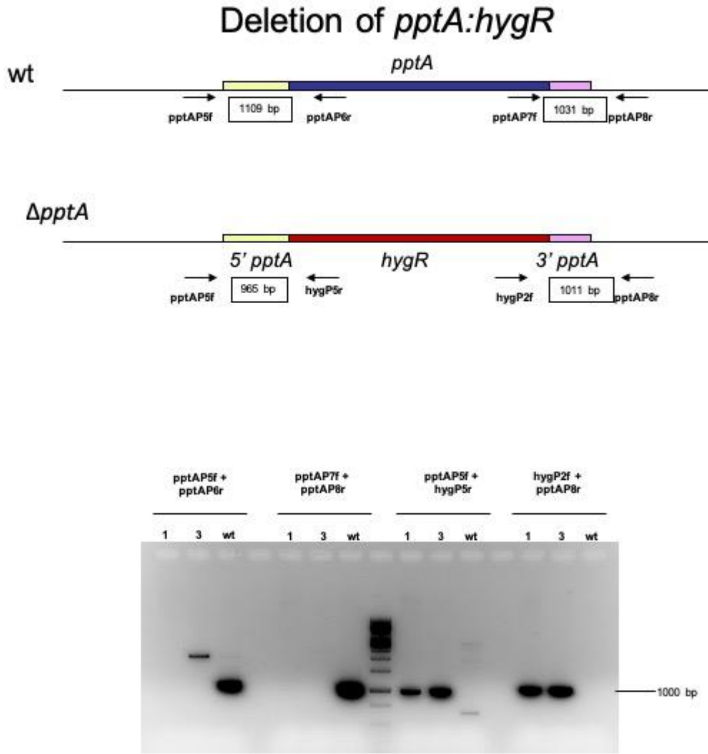

**Supplementary Figure 1.** Diagnostic PCR of the correct deletion of *pptA* in strain JN24.6, resulting in strain MA870.1. Clone 1 is used for experiments described. As a marker the GeneRuler 1 kb DNA ladder is used.

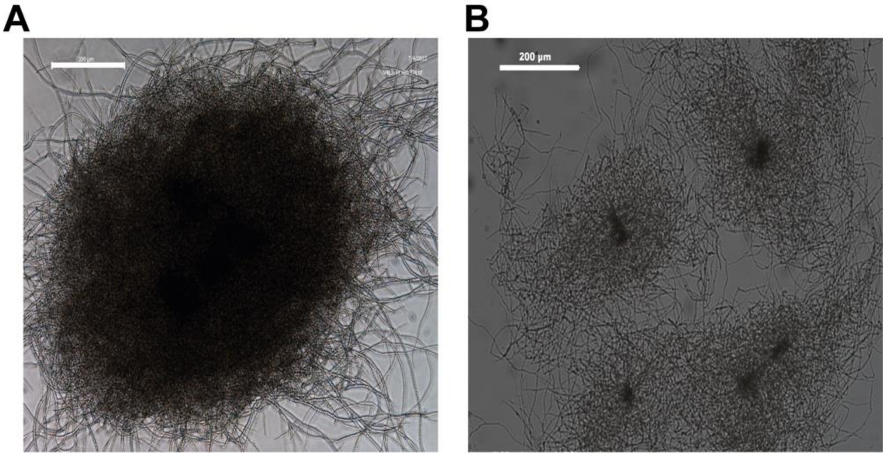

**Supplementary Figure 2.** Pellet formation of *A. niger* N402 (A) and *A. niger* D15#26 (B) after growing for 72 h  in MM supplemented with 25 mM xylose. Scale bare represents 200 µm.

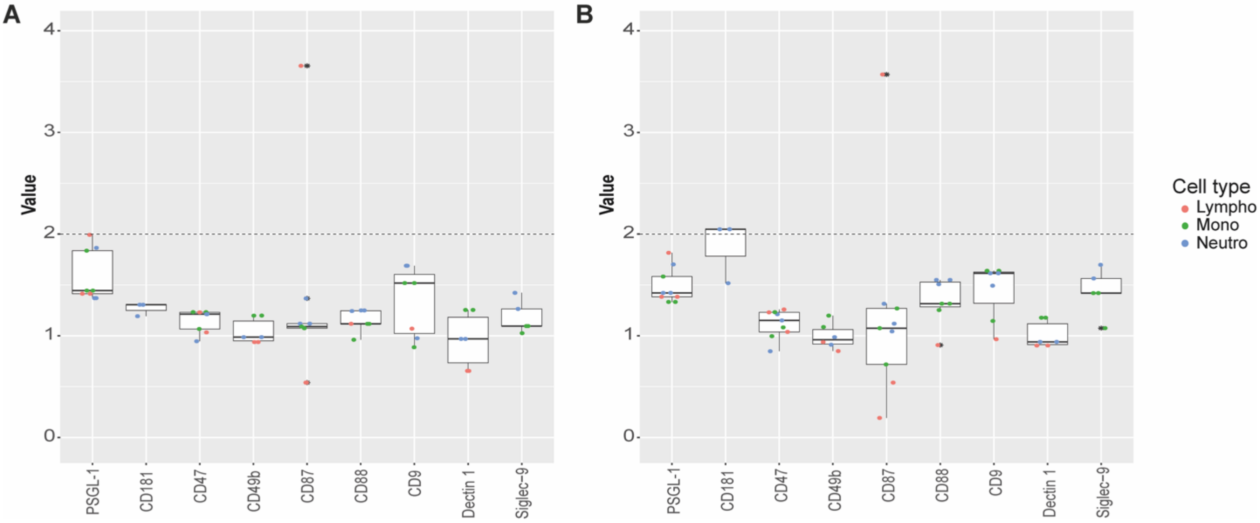

**Supplementary Figure 3**. Competition binding assay of D15#26 fractions after anion (A) or cation (B) chromatography. Lymphocytes, monocytes, and neutrophils are represented with red, green, and blue dots, respectively, * represent outliers. Y-axis represent the inverted geometric mean of fluorescence, the X-axis represent the used receptors in the CBA, data points above the dotted line (2) are scored as positive for binding of molecules from the culture medium. Samples were tested in 3 independent experiments.

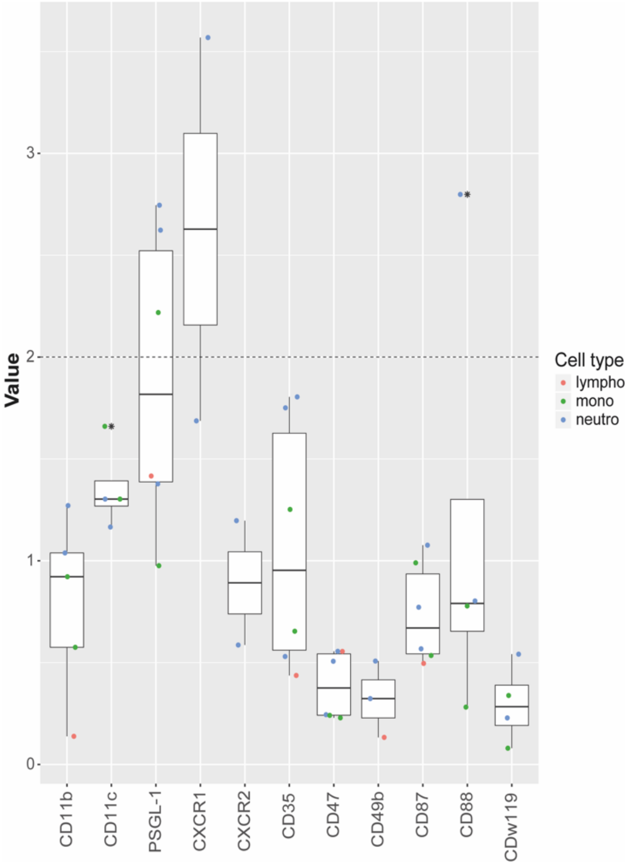

**Supplementary Figure 4.** Competition binding assay of 60 min – 100 °C treated D15#26 ≤ 3 kDa fractions. Lymphocytes, monocytes, and neutrophils are represented with red, green, and blue dots, respectively, * represent outliers. Y-axis represent the inverted geometric mean of fluorescence, the X-axis represent the used receptors in the CBA, data points above the dotted line (2) are scored as positive for binding of molecules from the culture medium. Samples were tested in 3 independent experiments.

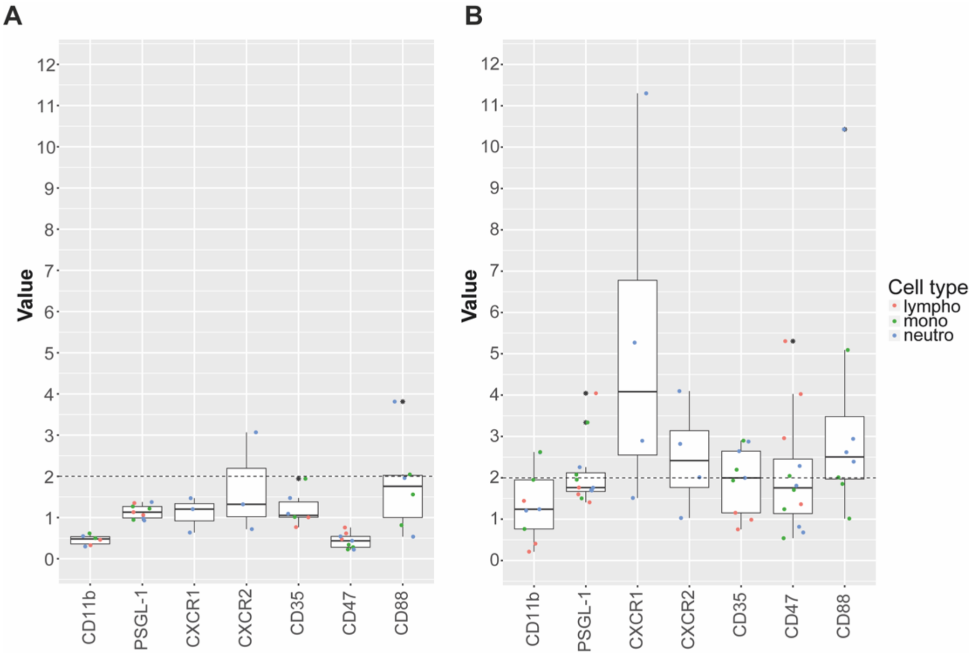

**Supplementary Figure 5.** Competition binding assay for D15#26 ≤ 3 kDa fractions after extraction with ethyl acetate (A) and the corresponding aqueous phase (B). Lymphocytes, monocytes, and neutrophils are represented with red, green, and blue dots, respectively, * represent outliers. Y-axis represent the inverted geometric mean of fluorescence, the X-axis represent the used receptors in the CBA, data points above the dotted line (2) are scored as positive for binding of molecules from the culture medium. Samples were tested in 3 independent experiments.

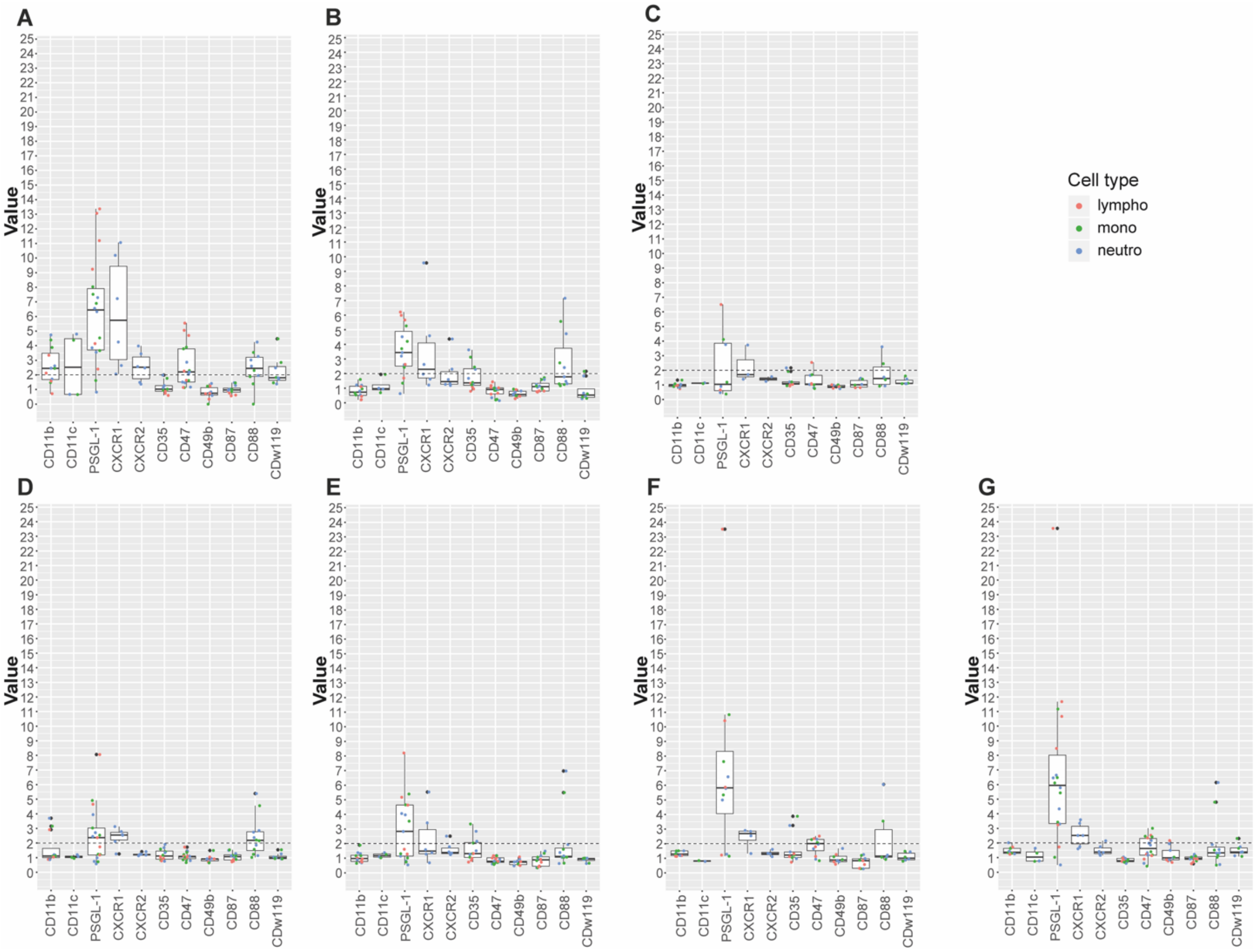

**Supplementary Figure 6**. Competition binding assay for D15#26 ≤ 3 kDa fractions after separation on a C18 Sep-pack column and elution with a gradient of methanol. Fractions used in the CBA: flowthrough (A), 0 % methanol (B), 10 % methanol (C), 30 % methanol (D), 50 % methanol (E), 70 % methanol (F) and 100 % methanol (G). Lymphocytes, monocytes, and neutrophils are represented with red, green, and blue dots, respectively, * represent outliers. Y-axis represent the inverted geometric mean of fluorescence, the X-axis represent the used receptors in the CBA, data points above the dotted line (2) are scored as positive for binding of molecules from the culture medium. Samples were tested in 3 independent experiments.

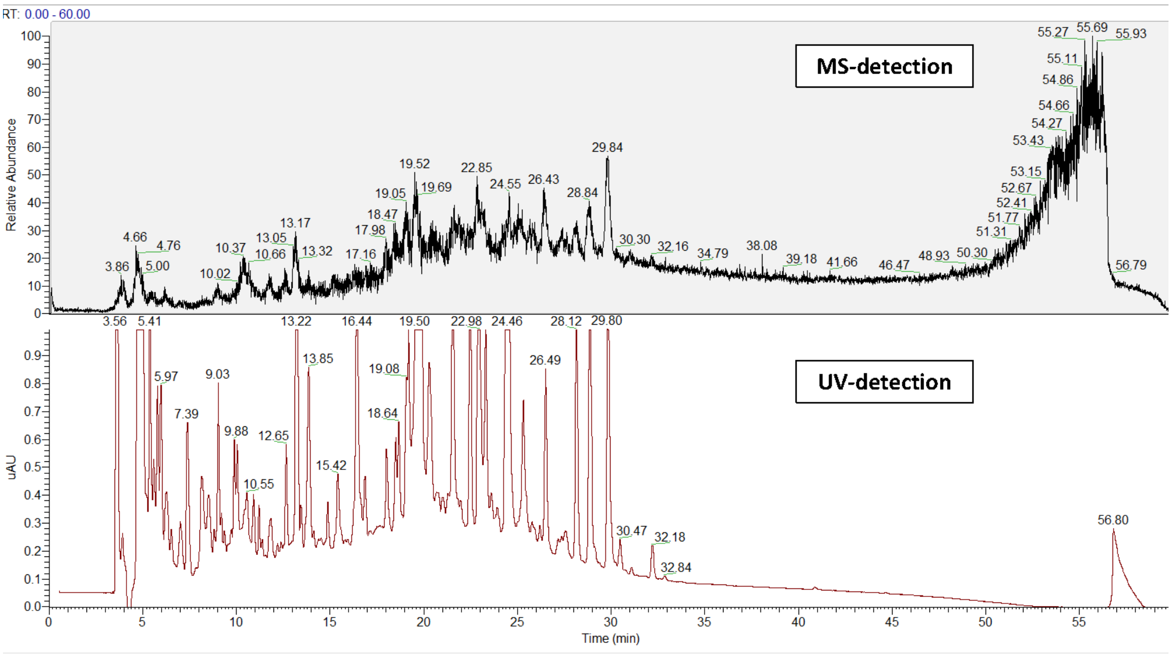

**Supplementary Figure 7.** LC-MS analysis of 10-50 % pooled methanol fractions after chromatography using a C-18 Sep-Pak column. Retention times were detected with MS detector and UV 214 nm.

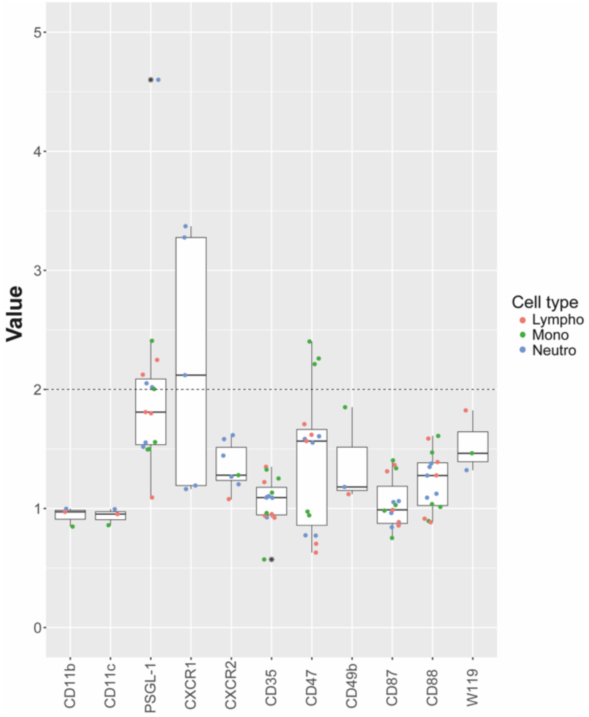

**Supplementary Figure 8.** Competition binding assay for *A. niger* AF11#56. Lymphocytes, monocytes, and neutrophils are represented with red, green, and blue dots, respectively, * represent outliers. Y-axis represent the inverted geometric mean of fluorescence, the X-axis represent the used receptors in the CBA, data points above the dotted line (2) are scored as positive for binding of molecules from the culture medium. Samples were tested in 3 independent experiments.

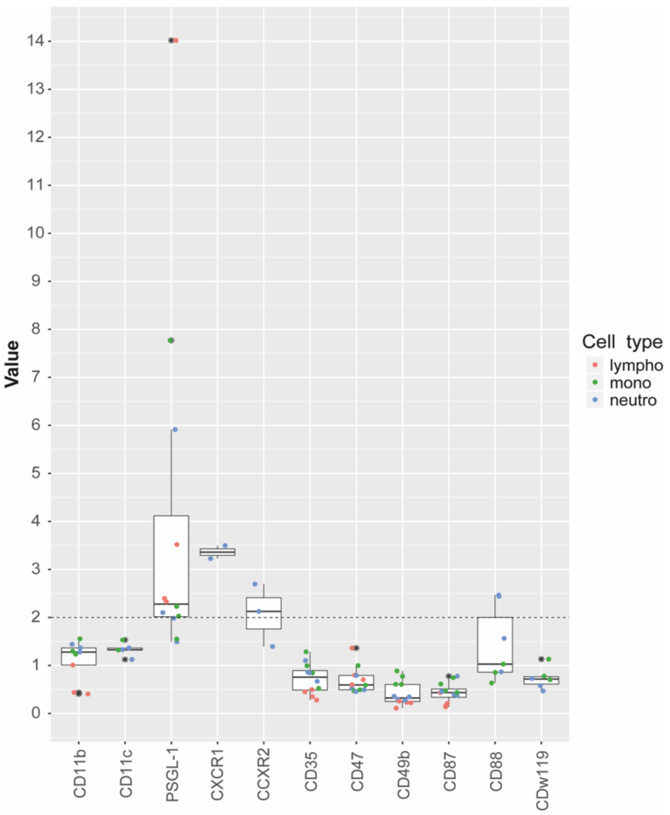

**Supplementary Figure 9.** Competition binding assay for *A. niger* Δ*crzA* ≤ 3 kDa fractions. Lymphocytes, monocytes, and neutrophils are represented with red, green, and blue dots, respectively, * represent outliers. Y-axis represent the inverted geometric mean of fluorescence, the X-axis represent the used receptors in the CBA, data points above the dotted line (2) are scored as positive for binding of molecules from the culture medium. Samples were tested in 3 independent experiments.

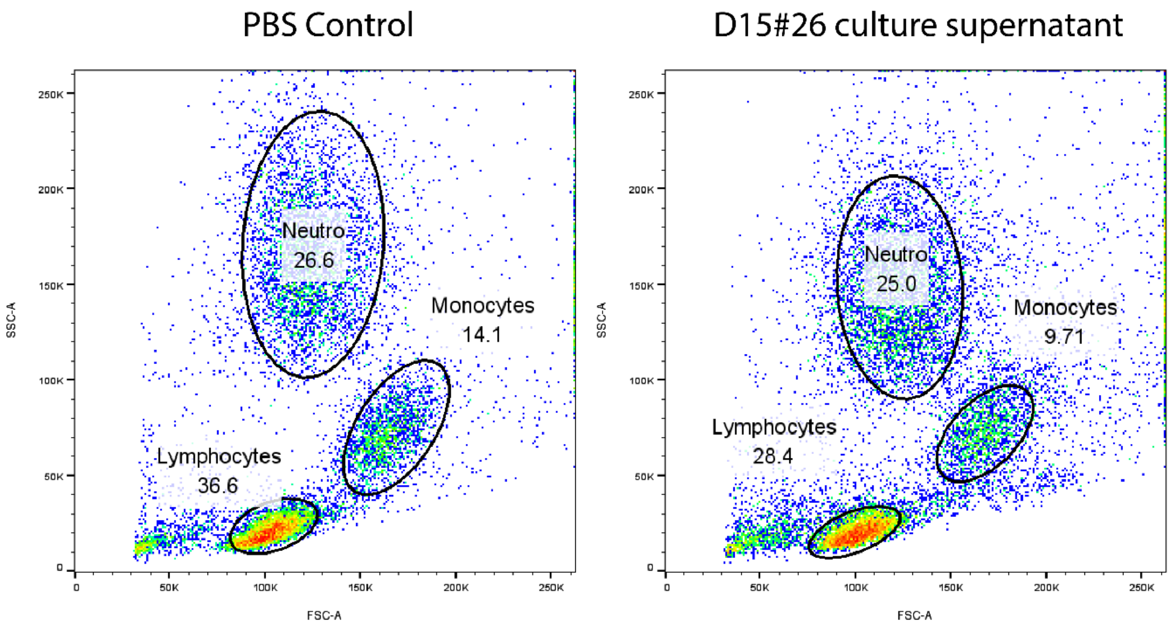

**Supplementary Figure 10**. Representative figure of the cell populations from the FACS analysis. Cells treated with a PBS control and D15#26 culture supernatant. No effects on intactness of cells was observed after treatment with fungal supernatants as determined in such FACS analysis.
